# Colloidally dispersed nanoparticles: a convergent evolutionary approach for the secretion of apparently soluble melanin

**DOI:** 10.64898/2026.02.06.704125

**Authors:** Senjuti Sarkar, Ananya Krishnaswamy, MB Govinda Rajulu, Pushya Pradeep, Shiwali Rana, Huma Sulthana, Pema Lhamu, Bhavana Chamraj, Deepika Rai, Rakshita S. Kolipakala, Judy Jays, N Thirunavukkarasu, JP Ravishankar, Sanjay K Singh, TS Suryanarayanan, Deepesh Nagarajan

## Abstract

Melanin is a heterogeneous, amorphous biopolymer widely distributed in biological systems, yet most canonical forms are intrinsically insoluble. Soluble melanins are rare and remain poorly characterized, with pyomelanin and peptidomelanin representing the best-described fungal examples. Here, we report the isolation and characterization of a distinct, polyphyletic class of soluble melanin that we term ‘colloidal melanin’. This class of polymers was isolated from four different species of Ascomycetous fungi occupying varied environmental niches. Colloidal melanin was found to be phenolic and nitrogenous, with a low amino acid content. The polymers form particles with median diameters ranging from 79 to 127 nm, suggesting that small particle size may contribute to aqueous solubility via colloidal dispersion. Using chemical inhibition of melanin biosynthesis together with whole-genome sequencing, we show that colloidal melanin can be produced through multiple distinct biochemical pathways. Further, genes identified as involved in melanin biosynthesis across species do not share any detectable common ancestry, consistent with convergent evolution. Together, these findings identify a distinct class of soluble melanin that does not conform to existing soluble melanin classifications.

## Introduction

Melanin is a structurally and chemically heterogeneous biopolymer widely distributed across biological kingdoms^1,2^. Although commonly recognized for its role in pigmentation, melanin represents a highly conserved class of biomolecules whose functions extend well beyond coloration^3–5^. Across diverse organisms, melanin contributes to photoprotection, free radical scavenging, metal ion chelation, and thermoregulation, reflecting its central role in environmental adaptation and cellular resilience^6,7^. The breadth of these functions underscores that melanin’s biological significance arises not merely from its optical properties, but from its underlying chemical composition^3,4,8^. Paradoxically, despite this functional versatility, most naturally occurring melanins are intrinsically insoluble in aqueous environments^1,9^, limiting their characterization using conventional techniques.

Chemically, melanins are amorphous macromolecules formed through the oxidative polymerization of phenolic or indolic precursors, including tyrosine / L-3,4-dihydroxyphenylalanine (L-DOPA)^10,11^, homogentisic acid (HGA) / gentisic acid^12,13^, and 1,8-dihydroxynaphthalene (DHN)^14,15^. Melanins can be broadly classified as eumelanin, pheomelanin, allomelanin, pyomelanin, and neuromelanin^8^. Eumelanins are derived from tyrosine and are synthesized along the Raper-Mason pathway^10,11^. Pheomelanin is a sulfur-containing (cysteinylated) polymer, producing reddish to yellow hues^16^. Neuromelanin occurs in the human brain^17,18^. Allomelanin is a nitrogen-free form derived from polyketide precursors like DHN^14^. Together, this biosynthetic and chemical diversity underpins the wide variation observed in melanin structure and behavior, including differences in molecular organization, reactivity, and solubility^19^.

Despite the extensive chemical and biosynthetic diversity observed among melanins, aqueous solubility remains uncommon. Within this context, attention has focused on a small number of naturally occurring soluble melanins. One of the best-characterized soluble melanins is pyomelanin, a nitrogen-free form derived from the polymerization of HGA during tyrosine catabolism^12^. Pyomelanin’s lower molecular weight and functional groups afford it high water solubility, allowing it to function as a secreted, redox-active protective agent in organisms like *Aspergillus fumigatus* and *Pseudomonas aeruginosa*^20^. Recent studies from our group identified a second water-soluble variant, peptidomelanin, produced by the fungus *Aspergillus niger* strain melanoliber^21,22^. Peptidomelanin is chemically more complex, consisting of an insoluble L-DOPA melanin core that is solubilized by short, covalently anchored, heterogeneous peptide chains^21,23,24^. Peptidomelanin is synthesized along the Srivatsan pathway^23^. Natural^21^ and synthetic^25^ forms of peptidomelanin can chelate heavy metals. This discovery established a new structural precedent: soluble melanin incorporating nitrogen. The presence of soluble melanin from other microbial species^9,26–30^ has been reported, however these forms have not been extensively biochemically characterized. The chemical synthesis of a soluble melanin with an increased abundance of carbonyl groups has also been reported^31^.

In this work, we identify and characterize soluble melanin secreted from four species of Ascomycetous fungi: *Gliomastix polychroma, Stachybotrys chlorohalonatus, Curvularia sp*., and *Pseudopithomyces sp*. We report that soluble melanin secreted from all four species is a nitrogenous, phenolic polymer. The nitrogen content suggests that the polymers are predominantly composed of L-DOPA, ruling out a predominantly pyomelanin composition. All polymers possessed a low or nil peptide content (below detection limit to 3.961 *±*0.671%), eliminating the possibility that the polymers are composed of peptidomelanin, and eliminating the possibility of nitrogen predominantly originating from a protein fraction. The polymers possess small particle diameters (79 to 127 nm); small even relative to other classes of melanin. This remains the most viable explanation for their apparent solubility: arising from colloidal dispersion rather than from the presence of solubilizing chemical groups.

Chemical inhibition of melanin synthesis using tropolone, tricyclazole, and 2-(2-nitro-4-trifluoromethylbenzoyl)-1,3-cyclohexanedione (NTBC), coupled with whole genome sequencing and functional annotation, revealed the involvement of different biochemical pathways producing colloidal melanin for every species. We detected involvement of the L-DOPA pathway for eumelanin, the 1,8-dihydroxynaphthalene (DHN) pathway for DHN melanin synthesis, and the homogentisic acid (HGA) pathway for pyomelanin synthesis. The involvement of these pathways for colloidal melanin synthesis differed across species. These results indicate that colloidal melanin is not an entirely new biochemical form of melanin. Rather, it can be created from existing melanin biosynthetic pathways, with minor modifications required to reduce particle size and impart solubility.

Whole genome sequencing and functional annotations revealed all genes involved in melanin biosynthesis across species. While we detected examples of paralogous genes, no orthologous melanin biosynthetic genes were detected. These observations, coupled with the differential involvement of known melanin biosynthetic pathways (as determined from melanin inhibition assays), indicate the convergent evolution of colloidal melanin.

We have chosen the term “colloidal melanin” to describe this class of polymers. Therefore, this work sits at the intersection of mycology and colloidal physics, which use different definitions for the term “soluble”. Mycology uses an operational definition: soluble melanin remains dispersed in water and resists removal by centrifugation or filtration. Colloidal physics reserves the term “soluble” for true solutions at the molecular scale, classifying larger particles (∼ 1–1000 nm) as colloidally dispersed rather than dissolved. It is therefore more accurate to describe our melanin polymers as “apparently soluble” (while continuing the use of the term “soluble” for brevity). This satisfies the mycological definition operationally, while also acknowledging colloidal dispersion and not true molecular dissolution.

## Results

### Fungal strain isolation

We isolated four filamentous fungi, sampled from diverse locations across South India and the Lakshadweep archipelago (Figure 1, Table 1). Fungi were isolated from diverse biomes and environmental media: urban garden soil, archipelagan coastal sponges, and from dry thorn forest leaf litter. The isolated species were identified as *Gliomastix polychroma* (VIG 6138 / NFCCI 6138), *Stachybotrys chlorohalonatus* (VIG 2586), *Curvularia sp*. (VIG 2862), and *Pseudopithomyces sp*. (VIG 728). The isolates were deposited into the Vivekananda Institute of Tropical Mycology (VINSTROM) group of cultures (VIG) or the National Fungal Culture Collection of India (NFCCI) for ease of access by the broader research community.

**Table 1.** Culture sampling data for all fungal cultures identified to secrete colloidal melanin.

| Species name | Culture ID | Collection date | Biome | Environmental medium | Location | Coordinates |
| --- | --- | --- | --- | --- | --- | --- |
| *Gliomastix polychroma* | VIG 6138 / NFCCI 6138 | 10<sup>th</sup> Jul 2008 | Urban / anthropogenic | Garden soil | RKM Vivekananda College campus, Chennai | 13.0408° N, 80.2644° E |
| *Stachybotrys chlorohalonatus* | VIG 2586 | 20<sup>th</sup> Dec 2022 | Coastal ocean | Sponge | Kavaratti island, Lakshadweep archipelago | 10.5593° N, 72.6358° E |
| *Curvularia sp.* | VIG 2862 | 17<sup>th</sup> May 2023 | Coastal ocean | Sponge | Agatti island, Lakshadweep archipelago | 10.8576° N, 72.1934° E |
| *Pseudopithomyces sp.* | VIG 728 | 18<sup>th</sup> Jul 2009 | Dry thorn forest | Leaf litter | Nilgiri Biosphere Reserve (NBR) | 11.53–11.72° N, 76.37°–76.75° E |

**Figure 1.**
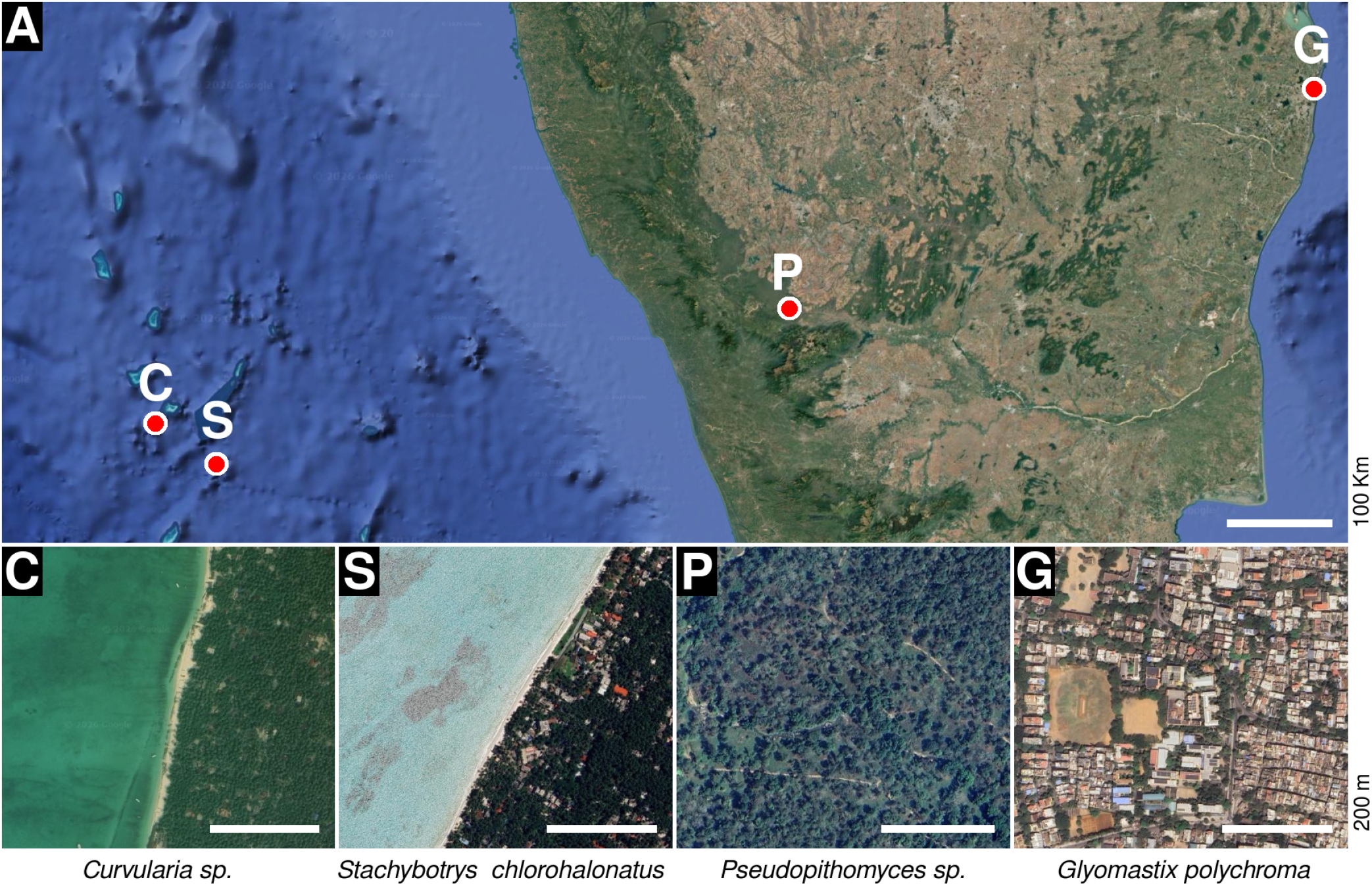
Satellite images (Google Earth) of the culture sampling locations of all colloidal melanin secreting strains described in this study. **(A)** The strains were sampled from across South India and the Lakshadweep archipelago. **(C)** *Curvularia sp*. was isolated from a sea sponge in the coast off Agatti island, Lakshadweep. **(S)** *Stachybotrys chlorohalonatus* was also isolated from a sea sponge in the coast off Kavaratti island, Lakshadweep. **(P)** *Pseudopithomyces sp*. was isolated from leaf litter in the Nilgiri Biosphere Reserve. **(G)** *Gliomastix polychroma* was isolated from garden soil in RKM Vivekananda College, Chennai. Detailed sampling data is provided in Table 1. For panel A, the scale bar represents 100 Km. For panels C,S,P, and G, the scale bar represents 200 m.

### Fungal strain identification and morphological characterization

The fungi were identified using molecular methods. For *Gliomastix polychroma* (VIG 6138 / NFCCI 6138), the internal transcribed spacer (ITS; GenBank accession ID: PX114506), large subunit ribosomal RNA gene (LSU; GenBank accession ID: PX114507), and translation elongation factor 1-alpha (EF-1*α*; GenBank accession ID: PX124606) gene regions were used to determine identity (Supplementary Text S1). For *Stachybotrys chlorohalonatus* (VIG 2586) the internal transcribed spacer (ITS; GenBank accession ID: PZ476833) was used to determine identity (Supplementary Text S2). For *Curvularia sp*. (VIG 2862) and *Pseudopithomyces sp*. (VIG 728), the internal transcribed spacer was extracted from their whole genome sequences (GenBank accession: JBZEGM000000000 and JBZEGL000000000, respectively) and used to determine identity. All ITS sequences are provided in Supplementary Text S3.

When grown on Sabouraud dextrose agar for 30 days, *Gliomastix polychroma* (VIG 6138 / NFCCI 6138) and *Stachybotrys chlorohalonatus* (VIG 2586) present as small, circular, melanated filamentous fungal colonies with dense aerial mycelia (Figures 2A and D, respectively). *Curvularia sp*. (VIG 2862) and *Pseudopithomyces sp*. (VIG 728) present as larger non-melanated filamentous fungal colonies with dense aerial mycelia (Figures 2G and J, respectively). Scanning electron microscopy reveals a dense, interconnected network of hyphae for all strains (Figures 2B-C,E-F,H-I,K-L). *Gliomastix polychroma* (VIG 6138 / NFCCI 6138) (Figure 2B,C) possessed densely-packed spores directly attached to the hyphal surface. No specialized conidia or spore-bearing structures were observed. No other strains displayed sporulation. It should be noted that sporulation is condition-dependent. The absence of spores under our growth conditions does not indicate the inability of the strains to sporulate altogether.

**Figure 2.**
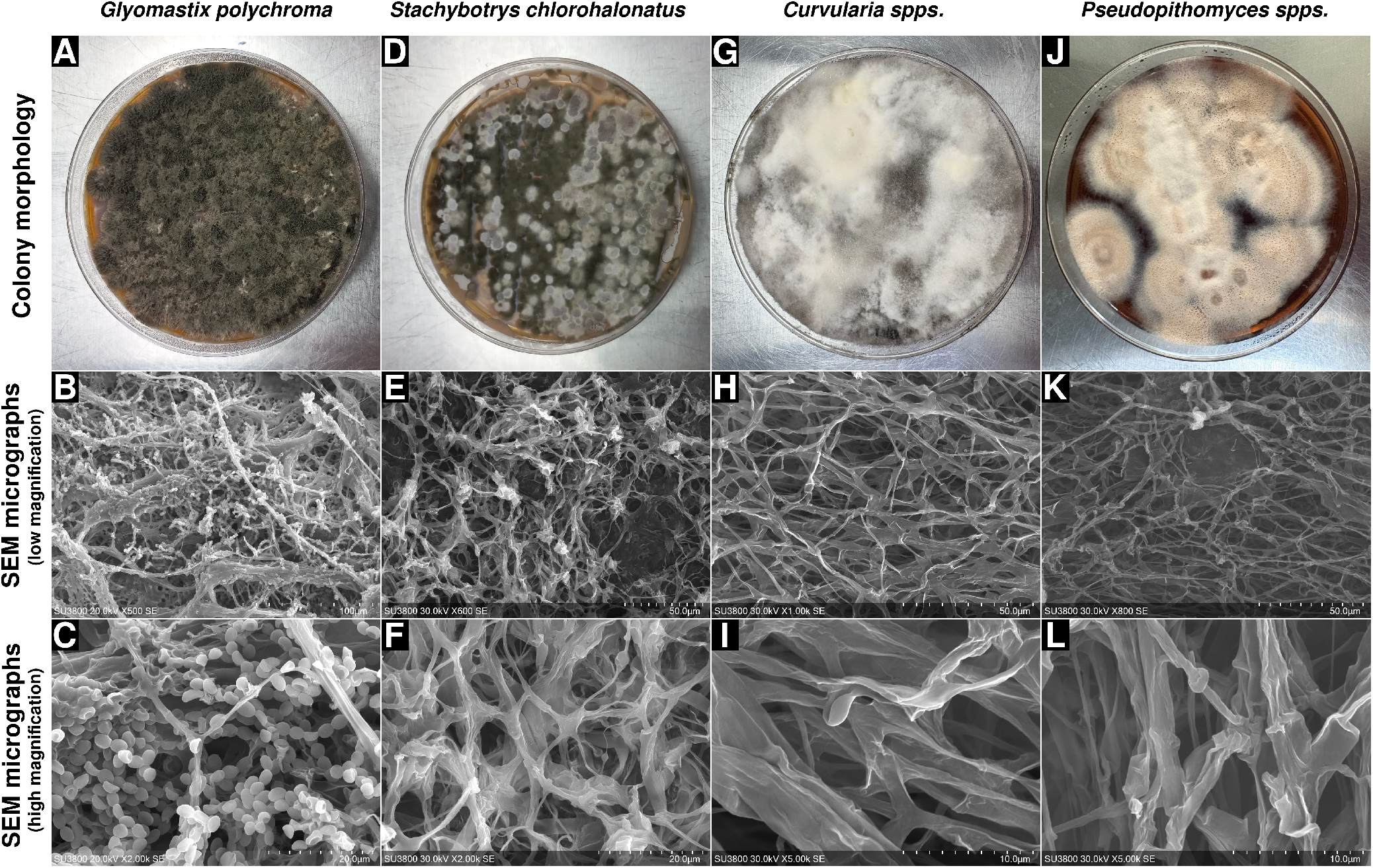
Strain morphology of all colloidal melanin-secreting strains. **(A–C)** *Gliomastix polychroma* (VIG 6138 / NFCCI 6138), **(D–F)** *Stachybotrys chlorohalonatus* (VIG 2586), **(G–I)** *Curvularia sp*. (VIG 2862), and **(J–L)** *Pseudopithomyces sp*. (VIG 728), showing (top row) colony morphology on agar after growth, (middle row) low-magnification and (bottom row) high-magnification scanning electron microscopy (SEM) micrographs of the mycelial network.

### Colloidal melanin conforms to the spectral characteristics of melanin standards

Colloidal melanin samples were extracted and purified as described in the methods section. The polymers were dissolved in deionized water, and appear as a black liquid (Figure 3A). All colloidal melanin samples display a UV-visible absorbance spectrum characteristic of melanin: broad absorption from 200–1000 nm (Figure 3B). Maximal absorbance was observed in the ultraviolet region (∼200 nm) with a continuous decrease in absorbance observed toward the near-infrared region (1000 nm), consistent with the optical behavior of melanin polymers. Similar spectra were also observed for synthetic L-DOPA melanin and natural peptidomelanin (from *A. niger* melanoliber)^21^.

**Figure 3.**
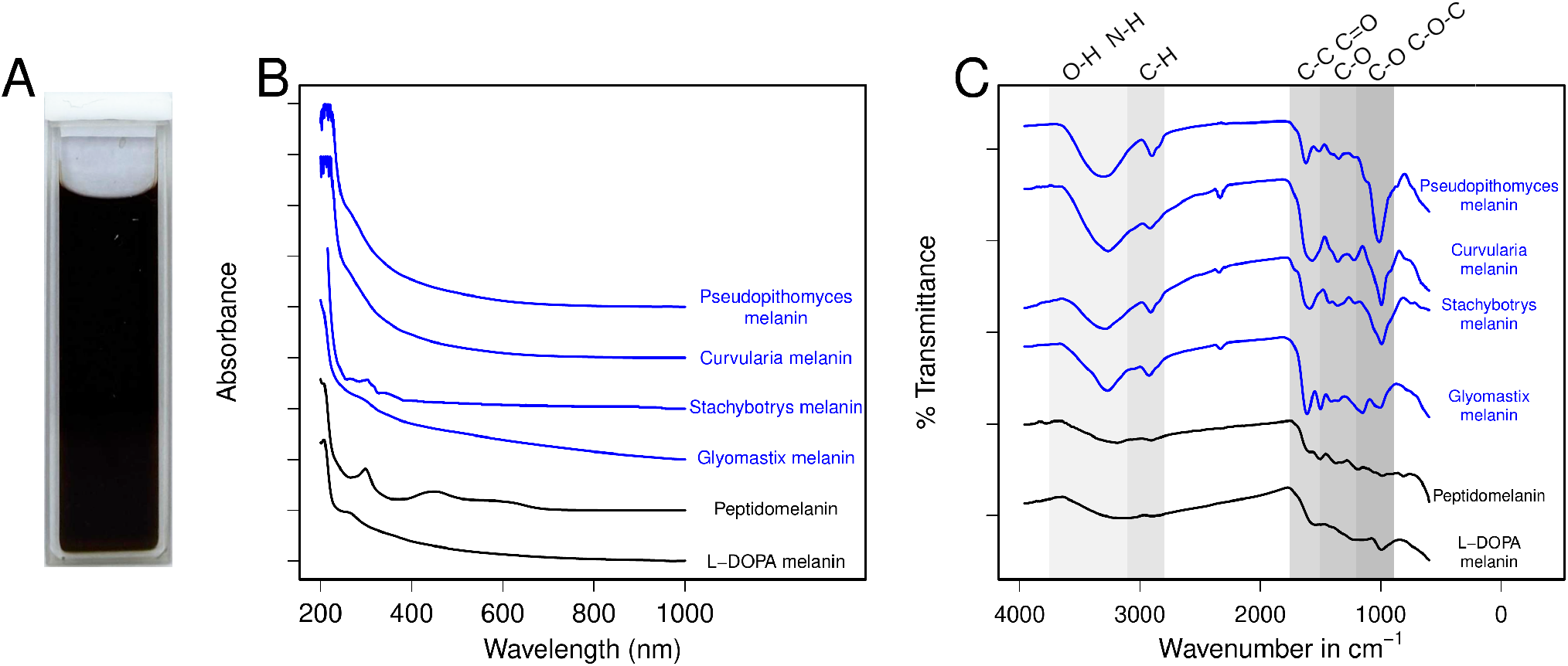
Spectroscopic characterization of colloidal melanins. **(A)** *Gliomastix polychroma* colloidal melanin (4 mg/mL in deionized water) observed in a quartz cuvette of 1 mm path length. The solution appears intensely black. All colloidal melanin samples appear identical to the naked eye. **(B)** The UV–visible absorption spectrum of colloidal melanin samples (0.2 mg/mL, blue) compared with reference melanins (peptidomelanin and L-DOPA melanin, 0.2 mg/mL, black). The spectra exhibits broad absorption from 200–1000 nm, with maximal absorbance in the ultraviolet region (∼200 nm) and a continuous decrease toward the near-infrared (1000 nm), consistent with the optical behavior of melanin polymers. Spectra have been vertically offset (by 0.2 a.u.) to facilitate comparison. **(C)** FTIR spectra of colloidal melanin samples (blue) and reference melanin samples (peptidomelanin and L-DOPA melanin, black) show broadly similar band distributions. All spectra have been vertically offset to ease comparison. The presence of dips within the 1500–1700 cm^−1^ region is attributable to overlapping aromatic ring vibrations and quinone-associated modes commonly reported for phenolic and indolic systems^5,7,21,32^.This supports the presence of phenolic, melanin-like chemical features.

The Fourier-transform infrared spectroscopy (FTIR) spectra of colloidal melanin samples were compared with that of natural peptidomelanin and synthetic L-DOPA melanin, revealing broadly similar absorption profiles characteristic of melanin pigments (Figure 3C). In particular, a prominent, overlapping dip spanning 1500–1700 cm^−1^ was observed, a region commonly associated with heterogeneous aromatic and quinone-rich structures in melanins. As expected for structurally heterogeneous biopolymers, these features do not permit unambiguous assignment of individual functional groups. Nevertheless, the overall spectral profile is consistent with established FTIR signatures reported for melanins^5,7,21,32^.

### Colloidal melanin is a heterogenous copolymer composed of monomers including L-DOPA, HGA, and DHN

To probe the biosynthetic processes responsible for synthesizing colloidal melanin across species, we performed melanin inhibition assays (Figure 4A,B) using tricyclazole, tropolone, and 2-(2-nitro-4-trifluoromethylbenzoyl)-1,3-cyclohexanedione (NTBC). All cultures, including inhibitor-treated samples and controls, were grown in medium supplemented with 0.5% tyrosine to ensure precursor availability. Treatment with tropolone and NTBC resulted in partial or complete colloidal melanin inhibition across all four strains, indicating a melanin polymer composed partly from L-DOPA and HGA (pyomelamin) monomers, respectively. In addition, tricyclazole partially or completely inhibited colloidal melanin production in *Gliomastix polychroma* (VIG 6138 / NFCCI 6138) and *Stachybotrys chlorohalonatus* (VIG 2586), indicating a melanin copolymer composed of L-DOPA, HGA (pyomelamin), and DHN monomers. These results indicate that, for a given species, colloidal melanin is synthesized through multiple biosynthetic routes rather than a single canonical pathway.

**Figure 4.**
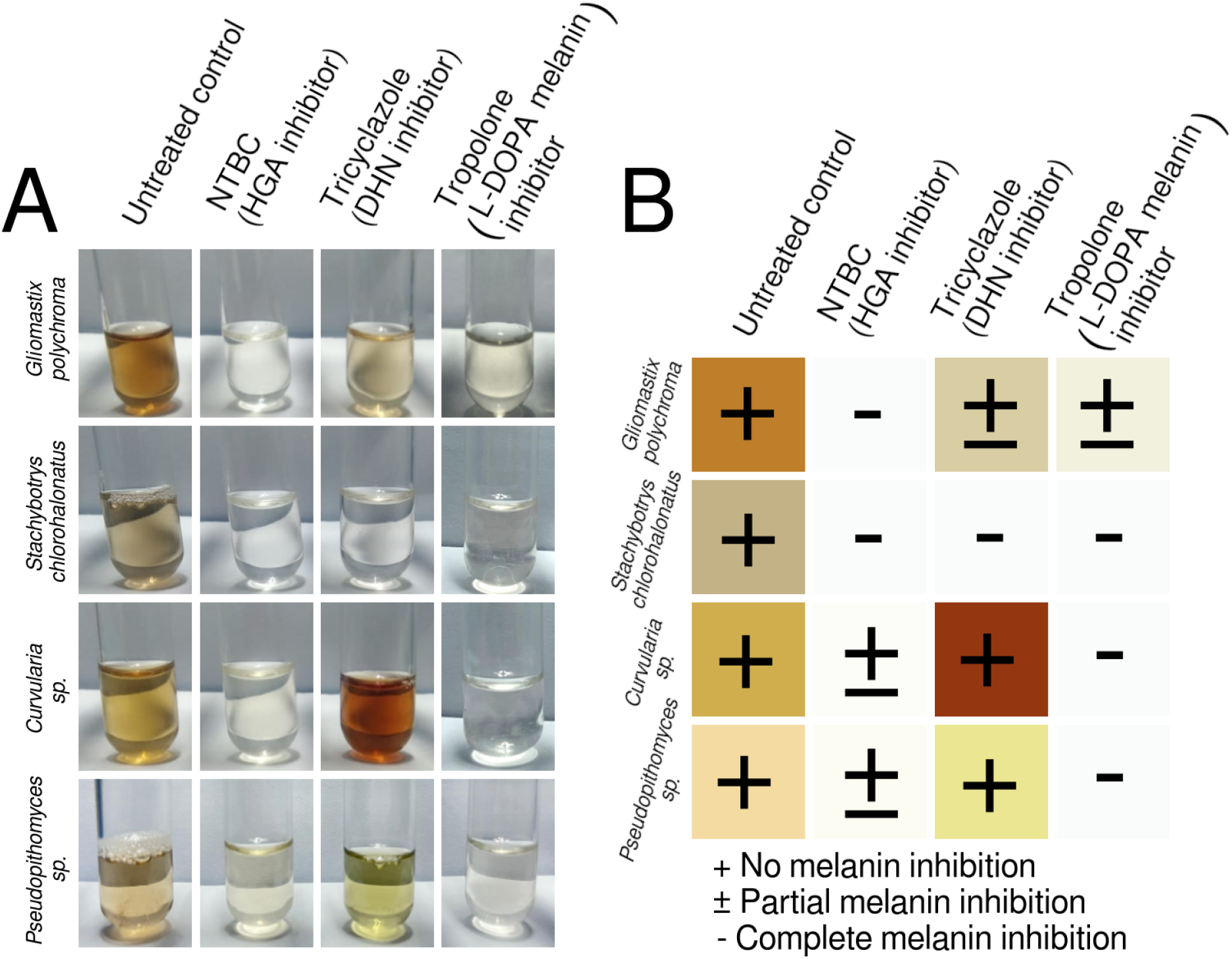
Melanin inhibition assays were performed using colloidal melanin secreting strains grown in Czapek Dox medium supplemented with 0.5% tyrosine. **(A)** Cultures grown under the following conditions: Untreated control; 1 mM NTBC, an inhibitor of 4-hydroxyphenylpyruvate dioxygenase (HPPD) in the pyomelanin (HGA melanin) pathway; 100 *µ*g/mL tricyclazole, which inhibits tetrahydroxynaphthalene reductase (THR) and 1,3,8-trihydroxynaphthalene reductase (1,3,8-THN) in the DHN-melanin pathway; and 2 mM tropolone, a well-established tyrosinase inhibitor, inhibiting the L-DOPA pathway. **(B)** Qualitative description of melanin inhibition (classified as none / partial / complete) for colloidal melanin secreting strains when treated with melanin inhibitors.

Taken together, the melanin inhibition profiles (Figure 4A,B) indicate that colloidal melanin production has evolved convergently across these four fungal strains rather than through a single conserved biosynthetic route. Each strain exhibited a distinct combination of pathway dependencies: *Gliomastix polychroma* (VIG 6138 / NFCCI 6138) required the HGA pathway obligately, with facultative contributions from the DHN and L-DOPA pathways. *Stachybotrys chlorohalonatus* (VIG 2586) required all three pathways obligately. *Curvularia sp*. (VIG 2862) and *Pseudopithomyces sp*. (VIG 728) required the L-DOPA pathway obligately, with facultative contributions from the HGA pathway. For these two strains, the DHN pathway was not required for the production of colloidal melanin. This absence of a shared inhibition profile across strains suggests that colloidal melanin secretion is not inherited from a common melanogenic ancestor but instead represents a convergent functional trait, independently assembled in each lineage from different combinations of the HGA, DHN, and L-DOPA biosynthetic pathways.

### Colloidal melanin possesses a significantly lower amino acid composition compared to peptidomelanin

Colloidal melanin is not a variant of naturally soluble peptidomelanin. Colloidal melanin samples were acid hydrolyzed to separate potential non-melanin conjugates from the melanin core polymer. The acid hydrolysate was subjected to liquid chromatography-mass spectrometry (LC-MS) to determine the amino acid content. The acid-hydrolyzed supernatant of all colloidal melanin samples exhibited significantly lower amino acid contents. The amino acid percentage (wt%) calculated for every colloidal melanin sample is as follows: *Gliomastix polychroma* (VIG 6138 / NFCCI 6138): 3.961 ± 0.671%; *Stachybotrys chlorohalonatus* (VIG 2586), 6.204 ± 0.774%; *Curvularia sp*. (VIG 2862), below detectable limit; and *Pseudopithomyces sp*. (VIG 728), below detectable limit. All these percentages are significantly lower compared to the reported 22.98 ± 1.84% amino acid content of peptidomelanin^21^ (Table 2).

**Table 2.**
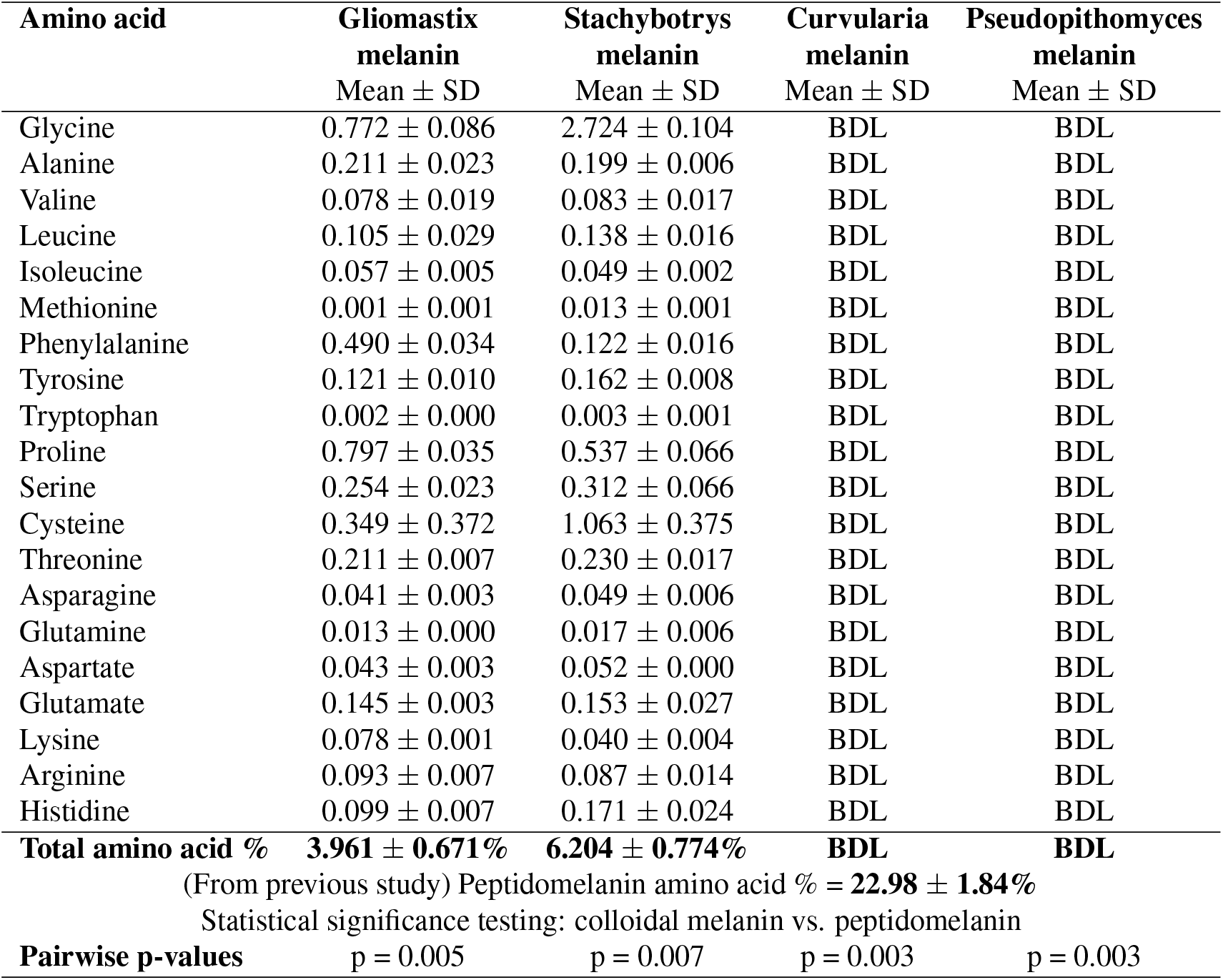
Liquid chromatography–mass spectrometry (LC–MS) of four acid-hydrolyzed colloidal melanin supernatants displayed low amino acid compositions (BDL to 6.204 ± 0.744%). In contrast to the 22.98 ± 1.84% amino acid content characteristic of peptidomelanin^21^, this significantly lower amino acid content (p = 0.003 to 0.007) indicates that the melanin polymers not composed of peptidomelanin. LC-MS reports and all calculations required to determine the final amino acid composition are provided in the supplementary information (**Dataset_S3:LC-MS_data**). BDL: Below detection limit.

### L-DOPA melanin comprises the largest single component of colloidal melanin

Colloidal melanin is a nitrogenous polymer. The elemental compositions of all colloidal melanin samples were determined using scanning electron microscopy - energy dispersive spectroscopy (SEM-EDS). These results are depicted in Table 3. The elemental analysis detected nitrogen, confirming the presence of an L-DOPA component of colloidal melanin previously predicted through tropolone inhibition assays (Figure 4). The nitrogen content of colloidal melanin samples, compared to L-DOPA melanin, is provided in Figure 5A. It should be noted that the nitrogen content (wt%) of colloidal melanin samples, although present, is significantly lower than that of chemically synthesized L-DOPA melanin. This can be explained by the copolymerization of non-nitrogenous melanin precursors (DHN and HGA) with nitrogenous L-DOPA melanin.

**Table 3.** Scanning electron microscopy - energy dispersive spectroscopy (SEM-EDS) elemental composition (C, N, O, S) of all melanin samples. Individual SEM-EDS spectra are provided in the supplementary information (**Dataset_S4:SEM-EDS_data**).

| Melanin polymer | C (wt%) | N (wt%) | O (wt%) | S (wt%) |
| --- | --- | --- | --- | --- |
| | Mean ± SD | Mean ± SD | Mean ± SD | Mean ± SD |
| L-DOPA melanin (synthetic) | 64.05 ± 1.47 | 10.19 ± 0.90 | 25.65 ± 0.87 | 0.11 ± 0.05 |
| Pyomelanin (synthetic) | 52.46 ± 5.62 | 0.00 ± 0.00 | 44.82 ± 4.76 | 2.71 ± 0.95 |
| Gliomastix melanin | 63.06 ± 3.31 | 7.58 ± 0.76 | 28.79 ± 2.65 | 0.57 ± 0.20 |
| Stachybotrys melanin | 73.63 ± 1.58 | 5.16 ± 0.54 | 21.01 ± 1.14 | 0.20 ± 0.06 |
| Curvularia melanin | 65.97 ± 9.03 | 4.97 ± 1.93 | 28.52 ± 10.28 | 0.53 ± 0.31 |
| Pseudopithomyces melanin | 66.77 ± 3.70 | 6.53 ± 1.16 | 26.17 ± 2.62 | 0.54 ± 0.21 |

**Figure 5.**
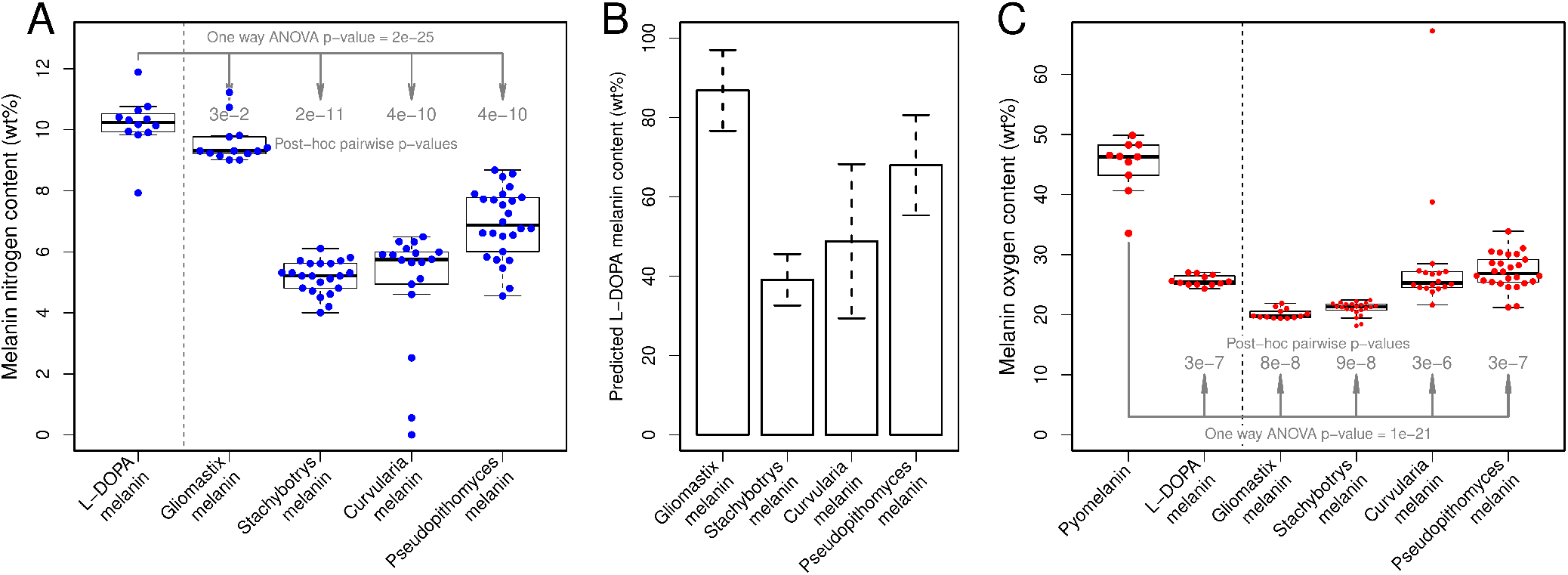
The elemental and molecular compositon of colloidal melanin. **(A)** Nitrogen content (wt%) of colloidal melanin compared to a chemically synthesized L-DOPA melanin standard. Nitrogen is present in all colloidal melanin samples, indicating a partial L-DOPA content. The nitrogen signal is significantly lower for colloidal melanins compared to L-DOPA melanin, indicating the presence of a non-nitrogenous melanin component. Blue dots represent individual data points. **(B)** L-DOPA component for all colloidal melanin samples, calculated from the nitrogenous component. The nitrogen signal arising from the peptide component of colloidal melanin (Table 2) was subtracted prior to this calculation. **(C)** Oxygen content of colloidal melanin compared to chemically synthesized L-DOPA melanin and pyomelanin standards. Colloidal melanin possesses an oxygen content (wt%) comparable to L-DOPA melanin, and significantly lower than pyomelanin, indicating that pyomelanin is not a major component of colloidal melanin. Red dots represent individual data points. Individual SEM-EDS spectra, along with R and bash scripts for statistical analysis and data visualization, are provided in the supplementary information (**Dataset_S4:SEM-EDS_data**).

L-DOPA melanin is the largest single component of colloidal melanin as a class of polymers. The percentage content of nitrogenous L-DOPA melanin (wt%) was directly quantified from the SEM-EDS nitrogen signal (Figure 5B). The nitrogen signal arising from the peptide component of colloidal melanin (Table 2) was subtracted prior to this calculation. The L-DOPA melanin percentage (wt%) calculated for every colloidal melanin sample is as follows: *Gliomastix polychroma* (VIG 6138 / NFCCI 6138): 86.85 ± 10.20%; *Stachybotrys chlorohalonatus* (VIG 2586), 39.08 ± 6.48%; *Curvularia sp*. (VIG 2862), 48.81 ± 19.44%; and *Pseudopithomyces sp*. (VIG 728), 68.00 ± 12.65%. Together, the mean percentage content of L-DOPA melanin across this class of polymers was found to be 60.69 ± 6.54%.

Colloidal melanin is not a variant of pheomelanin. Pheomelanin possesses a sulfur content of 6% to 16%^33^, far higher than the sulfur content (wt%) reported for colloidal melanin samples (0.20 *±* 0.06 to 0.57 *±* 0.20 (wt%), Table 3).

Pyomelanin, although present (Figure 4), is not a major component of colloidal melanin. Pyomelanin is a soluble melanin possessing a naturally elevated oxygen content^34–36^. We chemically synthesized pyomelanin from HGA, creating a polymer possessing an oxygen content (wt%) of 38.78 ± 2.91% (Table 3), significantly higher than the oxygen content (wt%) of chemically synthesized L-DOPA melanin or any of the colloidal melanin samples assayed (Figure 5C). These observations indicate that pyomelanin must be present as a minor component of colloidal melanin. It should be noted that unlike L-DOPA melanin, the pyomelanin content of colloidal melanin samples cannot be quantified as it possesses no unique elemental signal: the oxygen signal is shared across L-DOPA melanin, DHN melanin, and pyomelanin. The non-nitrogenous, non-pyomelanin (oxygen poor) component of colloidal melanin could potentially be composed of DHN melanin, although further experiments would be required to confirm its proportion.

Colloidal melanin is not solubilized due to oxygenous or nitrogenous functional groups. Pyomelanin, for example, is solubilized due to the presence of carboxyl and hydroxyl functional groups^37^. The absence of a strong oxygen signal for colloidal melanin (Figure 5C) eliminates the possibility of oxygen-rich functional groups of any origin imparting solubility to these polymers. Likewise, the absence of a strong nitrogen signal for colloidal melanin (Figure 5A) eliminates an alternative possibility of nitrogen-rich functional groups imparting solubility to these polymers.

### Evidence for solubility imparted through colloidal dispersion

In the previous sections, we have established that colloidal melanin is not solubilized due to its amino acid / peptide content (Table 2) or pyomelanin content (Figure 5C). We have also established that colloidal melanin lacks oxygenous or nitrogenous functional groups of any origin that could impart solubility (Figure 5C,A). Further, colloidal melanin is predominantly composed of insoluble L-DOPA melanin (Figure 5B), with the possibility of inclusion of large quantities of insoluble DHN melanin as well.

We therefore hypothesized that solubility may be the result of a biophysical process, rather than a biochemical process. Colloidal suspensions are created from small diameter nanoparticles that are insoluble in the true molecular sense, but appear soluble due to Brownian motion keeping them suspended in solution. As particle diameters grow, sedimentation forces come to dominate, forcing the suspension to precipitate.

We experimentally validated the hypothesis that L-DOPA melanin could be solubilized through colloidal suspension alone, without chemical modification. Firstly, we synthesized small-diameter L-DOPA melanin nanoparticles. The synthesis process used KMnO_4_ to rapidly oxidize L-DOPA in solution, limiting particle size. Small diameter L-DOPA melanin appeared soluble in aqueous solution (Figure 6A). Secondly, we synthesized large-diameter L-DOPA melanin nanoparticles. The synthesis process involved prolonged incubation at 80 ^°^C in DMSO with CuSO_4_ added as a catalyst for mild oxidation via atmospheric oxygen. Slow oxidation favors larger nanoparticles. In contrast, large-diameter L-DOPA melanin nanoparticles could not be suspended in aqueous solution (Figure 6C).

**Figure 6.**
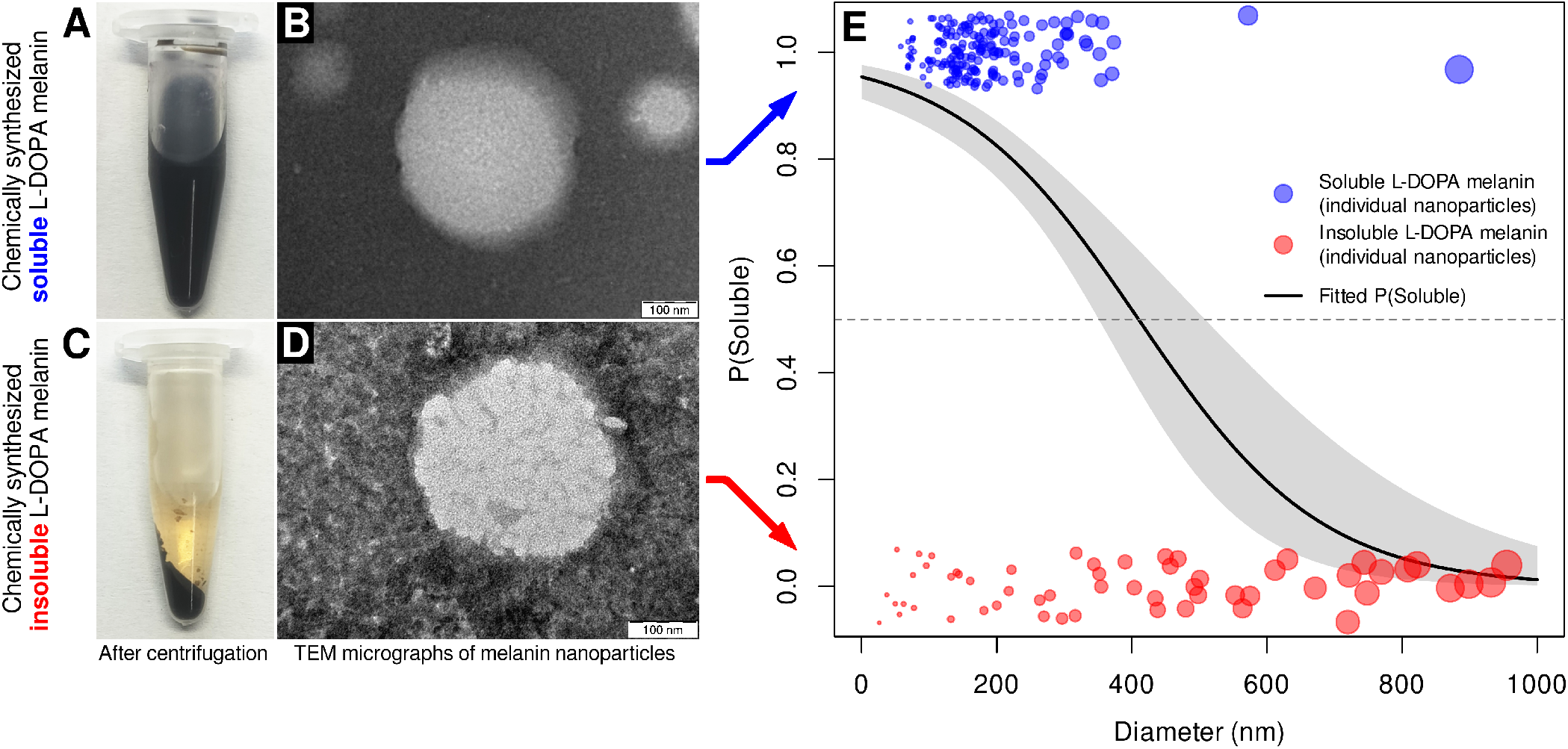
Chemical synthesis of soluble L-DOPA melanin (small-diameter) and insoluble L-DOPA melanin (large-diameter). **(A)** Soluble L-DOPA melanin, rapidly oxidized using KMnO_4_ to limit particle size, remains suspended after centrifugation at 7,000 rpm / 10 minutes. **(B)** Representative soluble L-DOPA melanin particle. 4 × digital zoom is applied. **(C)** Insoluble L-DOPA melanin, slowly oxidized using CuSO_4_ as a catalyst for atmospheric oxidation, appears in the pellet after centrifugation at 7,000 rpm / 10 minutes. **(D)** Representative insoluble L-DOPA melanin particle. **(B**,**D)** TEM micrographs depict representative particle morphologies (spherical). Particles appear white against phosphotungstic acid (negative stain). Particles shown are not representative of size. **(E)** Logistic regression model for the prediction of solubility based on nanoparticle diameter (*β* = −0.00739 ± 0.00123, *z* = −6.007, *p* = 1.89 × 10^−9^). Individual soluble / insoluble melanin nanoparticles are depicted in blue / red, respectively. Point sizes are proportional to nanoparticle diameters. The black line (fitted P(Soluble)) indicates predicted probability of solubility. The gray area represents 95% confidence intervals. Gray dotted line indicates P(Soluble) = 0.5 (50% probability of solubility threshold). All TEM micrographs along with R and bash scripts pertaining to data analysis, the logistic regression model, and data visualization, are provided in the supplementary information (**Dataset_S2:TEM_data**).

We confirmed nanoparticle synthesis using transmission electron microscopy (TEM). Both soluble and insoluble L-DOPA melanin were composed of spherical nanoparticles (Figure 6B,D). As expected, soluble L-DOPA melanin possessed smaller particle diameters (176 ± 90 nm) compared to insoluble L-DOPA melanin (397 ± 270 nm). Notably, insoluble L-DOPA melanin exhibited substantially greater variability in nanoparticle diameter. For this sample, larger diameter nanoparticles may inhibit the suspension of smaller diameter nanoparticles via sweep flocculation^38^.

We then trained a logistic regression model to classify the solubility of melanin nanoparticles based only on diameter (Figure 6E). Logistic regression confirmed a significant negative relationship between particle diameter and the probability of solubility (*β* = −0.00739 ± 0.00123, *z* = −6.007, *p* = 1.89 × 10^−9^). The model predicted a 50% probability of solubility at a diameter of 410 nm, with smaller particles predicted to remain soluble and larger particles predicted to be insoluble.

Transmission electron microscopy (TEM) revealed that colloidal melanin nanoparticles appear spherical (Figure 7A-D) and adopt size distributions compatible with colloidal dispersion. The mean diameters of colloidal melanin nanoparticles from *Gliomastix polychroma* (79 ± 36 nm), *Stachybotrys chlorohalonatus* (115 ± 59 nm), *Curvularia sp*. (127 ± 74 nm) and *Pseudopithomyces sp*. (84 ± 45 nm), are all smaller than that of synthetic soluble L-DOPA melanin (176 ± 90 nm) (Figure 7E). Further, the diameters of individual colloidal melanin nanoparticles are all classifiable as soluble (diameter *<*410 nm) using the logistic regression model trained on synthetic soluble and insoluble L-DOPA melanin (Figure 6). The same trends are observed when diameters are converted into volumes (Figure 7F): all colloidal melanin nanoparticles possess volumes classifiable as soluble using the same regression model. These results empirically demonstrate that, in the absence of detectable chemical functional groups to impart solubility, small particle diameters alone can solubilize both natural and synthetic melanin nanoparticles.

**Figure 7.**
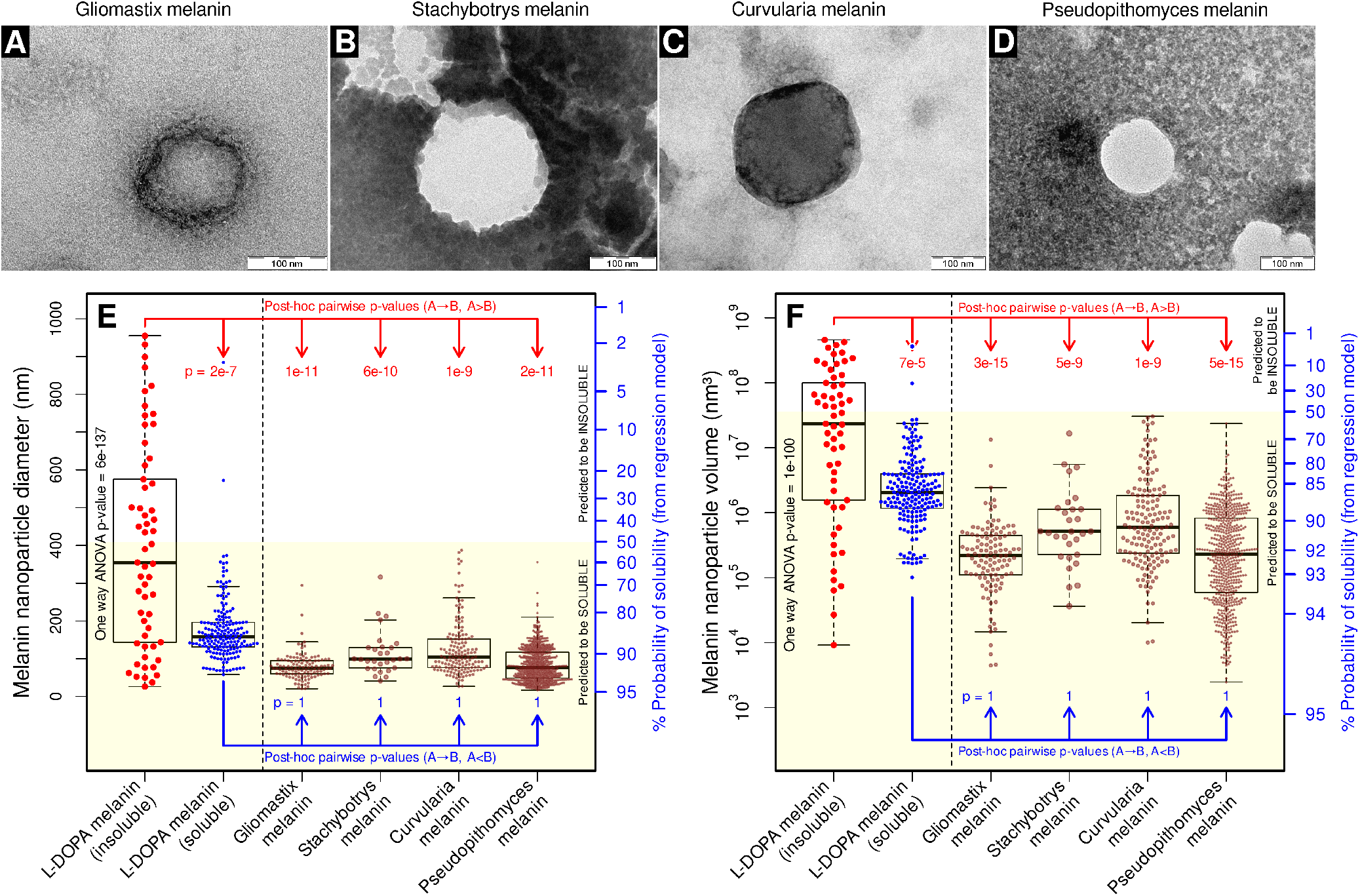
Colloidal melanin is composed of small diameter nanoparticles. Representative TEM micrographs of melanin nanoparticles secreted by **(A)** *Gliomastix polychroma* (VIG 6138 / NFCCI 6138), **(B)** *Stachybotrys chlorohalonatus* (VIG 2586), **(C)** *Curvularia sp*. (VIG 2862), and **(D)** *Pseudopithomyces sp*. (VIG 728). TEM micrographs depict representative particle morphologies (spherical). Particles were negatively stained using phosphotungstic acid (negative stain). Gliomastix melanin and curvularia melanin uptake the negative stain and appear darker. Particles shown are not representative of size. **(E)** Nanoparticle diameter distributions for insoluble L-DOPA melanin (red), soluble L-DOPA melanin (blue), and colloidal melanin samples (brown). Points represent the diameters of individual nanoparticles. All soluble melanin samples (soluble L-DOPA melanin and colloidal melanin samples) possessed particle diameters significantly smaller than insoluble L-DOPA melanin (red p-values). All colloidal melanin samples possessed particle diameters not significantly larger than soluble L-DOPA melanin (blue p-values). The yellow area highlights particle diameters predicted to display solubility by our logistic regression model. It encompasses nearly all particle diameters observed for soluble melanin samples (soluble L-DOPA melanin + colloidal melanin samples). **(F)** Our logistic regression model and statistical analyses produce the same predictions and same statistical inferences when particle volumes (rather than diameters) were analyzed. The volume corresponding to *P*(Soluble) = 0.5 (3.61 × 10^7^ nm^3^) was calculated directly from the corresponding diameter (410 nm). All TEM micrographs along with R and bash scripts pertaining to data analysis and visualization are provided in the supplementary information (**Dataset_S2:TEM_data**).

We complemented our empirical approach with a theoretical treatment of colloidal melanin dispersion. Within the field of colloidal physics, the relative influence of dispersive Brownian forces and sedimentation forces is modeled using the gravitational Péclet number (*Pe*_*g,c*_)^39^ (Equation 1), which scales with the fourth power of particle radius (*Pe*_*g,c*_ ∝ *r*^4^). Because larger *Pe*_*g,c*_ values indicate that sedimentation dominates over Brownian diffusion, colloidal suspendability is inversely proportional to particle radius to the fourth power.

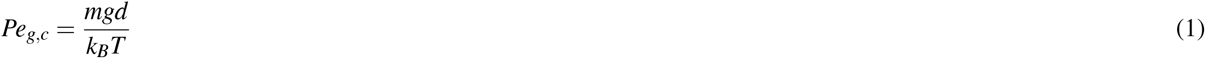

In Equation 1, *Pe*_*g,c*_ is the gravitational Péclet number, *m* is the buoyant mass of melanin (derived from melanin nanoparticle mean diameters, a melanin density of 1380 Kg/m^340^ and the density of water of 1000 Kg/m^3^), *d* is the mean diameter of melanin nanoparticles, *k*_*B*_ is the Boltzmann constant (1.380649 *×* 10^−23^ J/K), and T is the temperature (297.15 K).

The gravitational Péclet number of insoluble L-DOPA melanin was found to be 0.0551 (Table 4). As *Pe*_*g,c*_ ∝ *r*^4^, small decreases in diameter result in substantial increases in colloidal dispersion. The *Pe*_*g,c*_ of synthetic and natural colloidally dispersed melanin samples was 1-3 orders of magnitude (base 10) smaller than the *Pe*_*g,c*_ of insoluble L-DOPA melanin. *Pe*_*g,c*_ provides a theoretical framework to explain the colloidal dispersibility of melanin nanoparticles based only on their diameters.

**Table 4.** Mean diameters and gravitational Péclet numbers (Pe_*g,c*_) for colloidal melanin samples and synthetic controls.

| Sample | Diameter ($d$, nm)<br>(mean ± SD) | Diameter ratio =<br>$\frac{d_{\text{Sample}}}{d_{\text{L-DOPA melanin (insoluble)}}}$ | Gravitational Péclet number<br>($Pe_{g,c}$) | Péclet ratio =<br>$\frac{Pe_{g,c} \text{ Sample}}{Pe_{g,c} \text{ L-DOPA melanin (insoluble)}}$ |
| --- | --- | --- | --- | --- |
| L-DOPA melanin (insoluble) | 397 ± 270 | 1.00× | $5.51 \times 10^{-2}$ | 1× |
| L-DOPA melanin (soluble) | 176 ± 90 | 0.44× | $2.75 \times 10^{-3}$ | 20× |
| Gliomastix melanin | 79 ± 36 | 0.20× | $6.86 \times 10^{-5}$ | 804× |
| Stachybotrys melanin | 115 ± 59 | 0.29× | $3.16 \times 10^{-4}$ | 175× |
| Curvularia melanin | 127 ± 74 | 0.32× | $6.06 \times 10^{-4}$ | 91× |
| Pithomyces melanin | 84 ± 45 | 0.21× | $8.97 \times 10^{-5}$ | 615× |

### Comparative genomics supports the convergent evolution of colloidal melanin

We performed whole-genome sequencing (WGS) of the four fungal strains that secrete colloidal melanin. Genome accession numbers for the genome of each strain are as follows: *Gliomastix polychroma* (VIG 6138 / NFCCI 6138): JBSVRS000000000.1; *Stachybotrys chlorohalonatus* (VIG 2586): JBYLTQ000000000; *Curvularia sp*. (VIG 2862): JBZEGM000000000; and *Pseudopithomyces sp*. (VIG 728): JBZEGL000000000. The functional annotation of the WGS data revealed genes involved in melanin synthesis through the L-DOPA, DHN, and HGA pathways (Table 5), supporting our observations from melanin inhibitor assays (Figure 4).

**Table 5.**
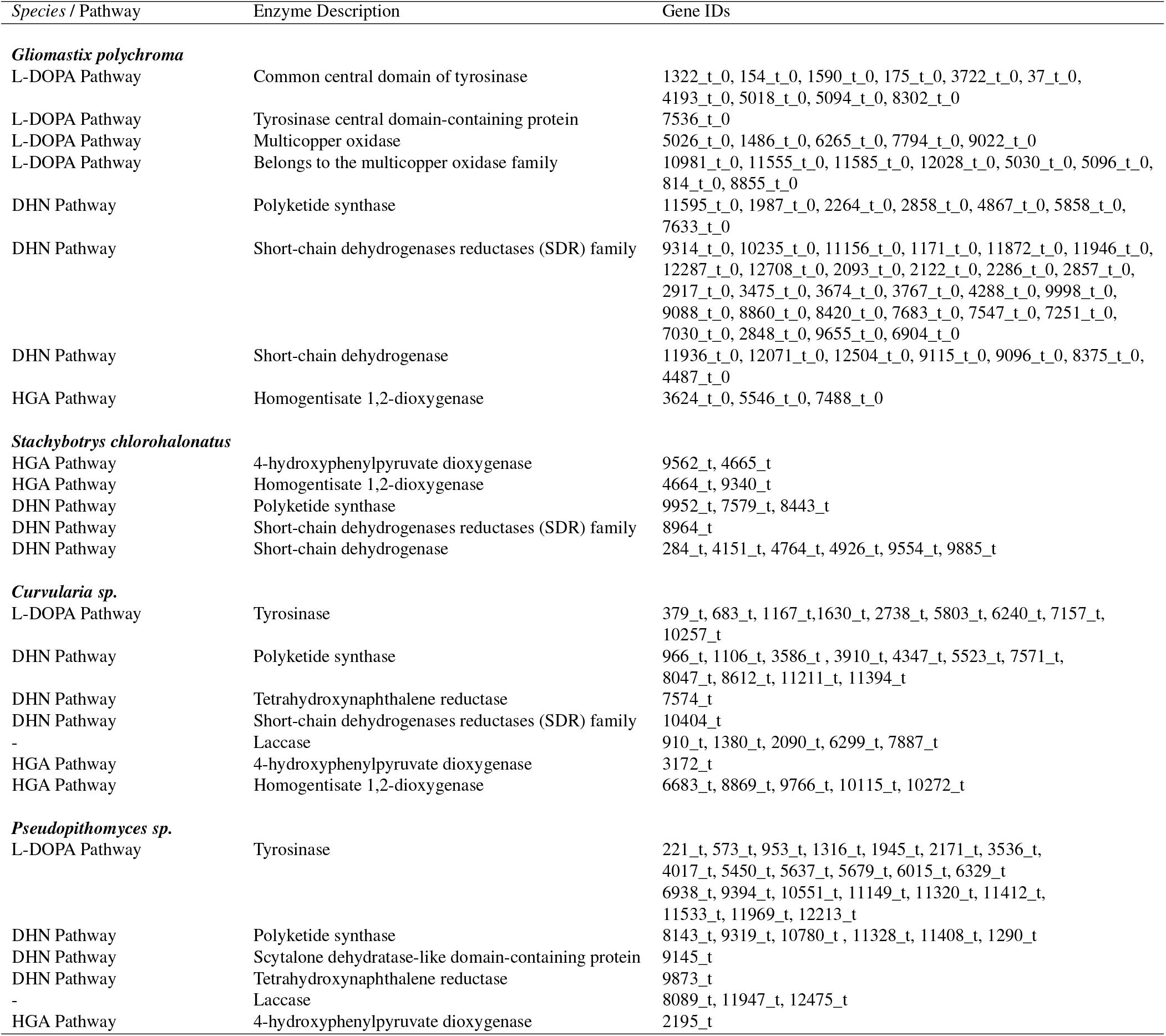
Whole-genome sequencing–based identification of genes involved in melanin biosynthesis from all four strains observed to secrete colloidal melanin. Annotated genes corresponding to key enzymatic steps in established fungal melanin biosynthetic pathways are listed. Detailed annotations for genes involved in melanin synthesis for every strain are provided in the supplementary information (Tables S1-S4). Gene sequences for every gene involved in melanin synthesis is provided in the supplementary information (**Dataset S5:annotated_gene_sequences**).

| Species / Pathway | Enzyme Description | Gene IDs |
| --- | --- | --- |
| ***Glomastix polychroma*** |  |  |
| L-DOPA Pathway | Common central domain of tyrosinase | 1322_t_0, 154_t_0, 1590_t_0, 175_t_0, 3722_t_0, 37_t_0, 4193_t_0, 5018_t_0, 5094_t_0, 8302_t_0 |
| L-DOPA Pathway | Tyrosinase central domain-containing protein | 7536_t_0 |
| L-DOPA Pathway | Multicopper oxidase | 5026_t_0, 1486_t_0, 6265_t_0, 7794_t_0, 9022_t_0 |
| L-DOPA Pathway | Belongs to the multicopper oxidase family | 10981_t_0, 11555_t_0, 11585_t_0, 12028_t_0, 5030_t_0, 5096_t_0, 814_t_0, 8855_t_0 |
| DHN Pathway | Polyketide synthase | 11595_t_0, 1987_t_0, 2264_t_0, 2858_t_0, 4867_t_0, 5858_t_0, 7633_t_0 |
| DHN Pathway | Short-chain dehydrogenases reductases (SDR) family | 9314_t_0, 10235_t_0, 11156_t_0, 1171_t_0, 11872_t_0, 11946_t_0, 12287_t_0, 12708_t_0, 2093_t_0, 2122_t_0, 2286_t_0, 2857_t_0, 2917_t_0, 3475_t_0, 3674_t_0, 3767_t_0, 4288_t_0, 9998_t_0, 9088_t_0, 8860_t_0, 8420_t_0, 7683_t_0, 7547_t_0, 7251_t_0, 7030_t_0, 2848_t_0, 9655_t_0, 6904_t_0 |
| DHN Pathway | Short-chain dehydrogenase | 11936_t_0, 12071_t_0, 12504_t_0, 9115_t_0, 9096_t_0, 8375_t_0, 4487_t_0 |
| HGA Pathway | Homogentisate 1,2-dioxygenase | 3624_t_0, 5546_t_0, 7488_t_0 |
| ***Stachybotrys chlorohalonatus*** |  |  |
| HGA Pathway | 4-hydroxyphenylpyruvate dioxygenase | 9562_t, 4665_t |
| HGA Pathway | Homogentisate 1,2-dioxygenase | 4664_t, 9340_t |
| DHN Pathway | Polyketide synthase | 9952_t, 7579_t, 8443_t |
| DHN Pathway | Short-chain dehydrogenases reductases (SDR) family | 8964_t |
| DHN Pathway | Short-chain dehydrogenase | 284_t, 4151_t, 4764_t, 4926_t, 9554_t, 9885_t |
| ***Curvularia sp.*** |  |  |
| L-DOPA Pathway | Tyrosinase | 379_t, 683_t, 1167_t, 1630_t, 2738_t, 5803_t, 6240_t, 7157_t, 10257_t |
| DHN Pathway | Polyketide synthase | 966_t, 1106_t, 3586_t, 3910_t, 4347_t, 5523_t, 7571_t, 8047_t, 8612_t, 11211_t, 11394_t |
| DHN Pathway | Tetrahydroxynaphthalene reductase | 7574_t |
| DHN Pathway | Short-chain dehydrogenases reductases (SDR) family | 10404_t |
| - | Laccase | 910_t, 1380_t, 2090_t, 6299_t, 7887_t |
| HGA Pathway | 4-hydroxyphenylpyruvate dioxygenase | 3172_t |
| HGA Pathway | Homogentisate 1,2-dioxygenase | 6683_t, 8869_t, 9766_t, 10115_t, 10272_t |
| ***Pseudopithomyces sp.*** |  |  |
| L-DOPA Pathway | Tyrosinase | 221_t, 573_t, 953_t, 1316_t, 1945_t, 2171_t, 3536_t, 4017_t, 5450_t, 5637_t, 5679_t, 6015_t, 6329_t, 6938_t, 9394_t, 10551_t, 11149_t, 11320_t, 11412_t, 11533_t, 11969_t, 12213_t |
| DHN Pathway | Polyketide synthase | 8143_t, 9319_t, 10780_t, 11328_t, 11408_t, 1290_t |
| DHN Pathway | Scytalone dehydratase-like domain-containing protein | 9145_t |
| DHN Pathway | Tetrahydroxynaphthalene reductase | 9873_t |
| - | Laccase | 8089_t, 11947_t, 12475_t |
| HGA Pathway | 4-hydroxyphenylpyruvate dioxygenase | 2195_t |

It should be noted that no genes involved in the L-DOPA pathway were detected in *Stachybotrys chlorohalonatus* (VIG 2586), despite the presence of nitrogen (Figure 5A,B) and inhibition by tropolone (Figure 4) indicating the presence of L-DOPA melanin. It should be noted that fungi possess a large number of tyrosinase isozymes. It is therefore possible that undiscovered (and therefore unannotated) tyrosinase sequences may exist within the *Stachybotrys chlorohalonatus* genome.

Additionally, *Curvularia sp*. (VIG 2862) and *Pseudopithomyces sp*. (VIG 728) possessed genes involved in the DHN pathway, despite displaying no colloidal melanin inhibition when treated with tricyclazole (DHN melanin inhibitor, Figure 4). For these two species, we can infer that DHN melanin is not an essential component of their colloidal melanin polymers.

The previously-described melanin inhibition assays (Figure 4A,B) showed that each strain possessed distinct pathways for colloidal melanin synthesis; a strong indication of convergent evolution. Supporting this conclusion, we detected no orthologous melanin biosynthetic genes across all four fungal strains (Figure 8A,B). Figure 8A depicts a comparison of pairwise sequence alignment identity (%) scores for unshuffled and shuffled melanin biosynthetic genes. From Table 5, we identified 150 melanin biosynthetic genes from all four strains (69 from *Gliomastix polychroma* (VIG 6138 / NFCCI 6138), 14 from *Stachybotrys chlorohalonatus* (VIG 2586), 33 from *Curvularia sp*. (VIG 2862), 34 from *Pseudopithomyces sp*. (VIG 728)).

**Figure 8.**
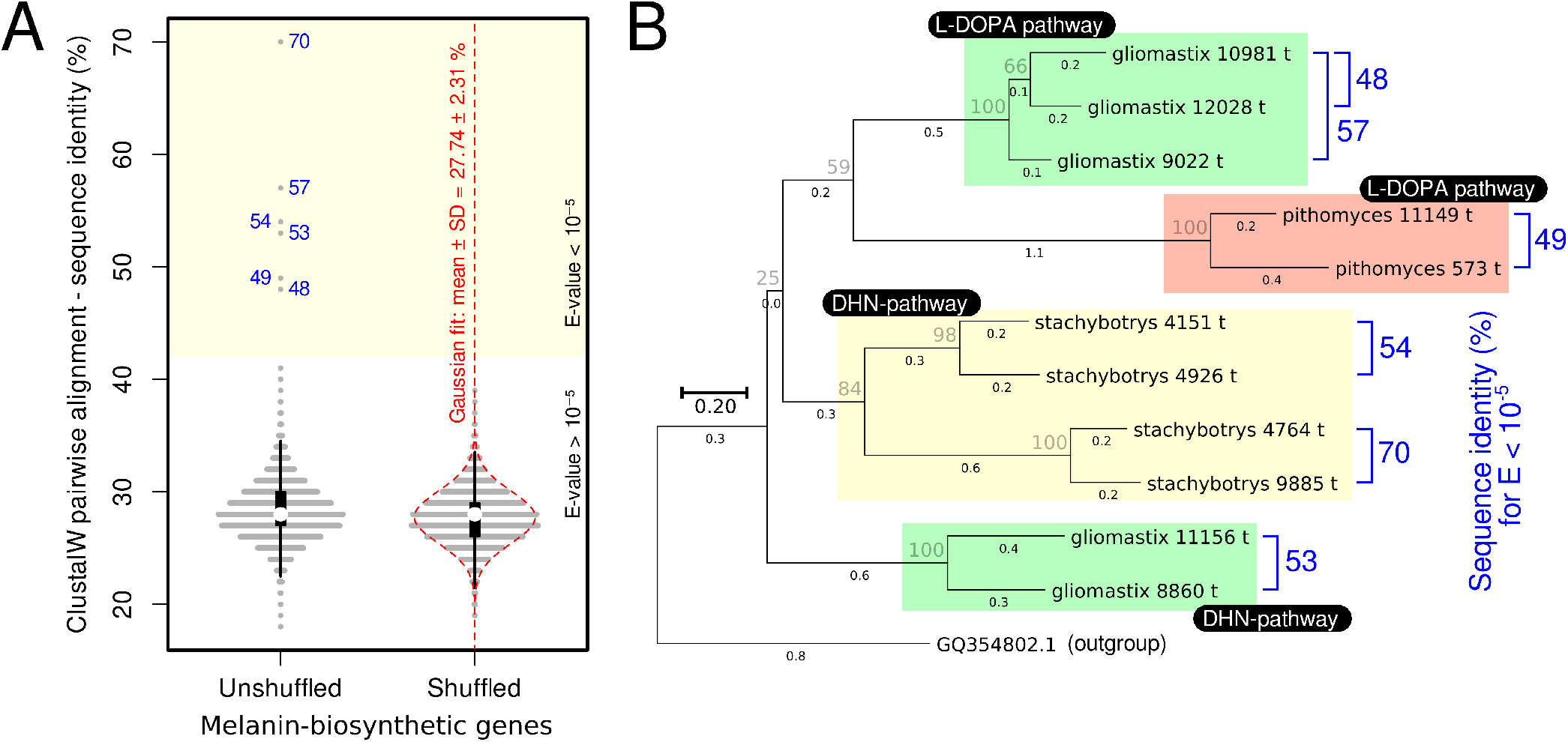
Paralogous, but not orthologous, melanin biosynthetic genes identified within colloidal melanin secreting fungal strains. **(A)** For unshuffled: ClustalW pairwise, non-redundant gene alignments of 150 melanin biosynthetic genes were annotated from the whole genome sequences of four fungal strains (150C_2_ = 11175 pairwise comparisons). Gray horizontal bars represent frequency distributions of pairwise sequence alignment identities. Boxplots (in black) are overlaid. Yellow region represents pairwise sequence alignment identities ≥42% (E ≤ 10^−5^). 6 pairwise sequence alignment identities (blue numbers) exist beyond this significance threshold. For shuffled: The sequences of 150 melanin biosynthetic genes were shuffled while keeping their codons intact. 150C_2_ = 11175 ClustalW pairwise, non-redundant gene alignments were then performed as for unshuffled genes. Red dotted line indicates Gaussian fit. The pairwise alignment score (42%) corresponding to an E-value of 10^−5^ was derived from this dataset. **(B)** Maximum parsimony phylogenetic tree (100 bootstrap replicates) constructed from melanin biosynthetic gene sequences possessing ≥42% (E ≤ 10^−5^) pairwise sequence alignment identities. The phylogenetic tree establishes the presence of paralogous genes, but not orthologous genes. Gray values indicate bootstrap support values. Black values indicate branch lengths. The scale bar (0.20) denotes the scale of branch lengths.

Firstly, we performed ClustalW pairwise, non-redundant gene alignments for all 150 melanin biosynthetic genes (unshuffled) (150C_2_ = 11175 pairwise comparisons). The distribution of pairwise sequence alignment identities (%) is provided in Figure 8A, unshuffled. This distribution possessed a mean pairwise sequence alignment identity of 28.20 *±* 2.53 (%).

Secondly, sequences of 150 melanin biosynthetic genes were randomly shuffled while keeping the individual codons intact (Figure 8, shuffled), before performing ClustalW pairwise, non-redundant gene alignments (150C_2_ = 11175 pairwise comparisons). This distribution possessed a mean pairwise sequence alignment identity of 27.74 ± 2.31 %, and fit a Gaussian distribution. This distribution represents the expected observations for pairwise sequence alignment identity (%) for a dataset of genes unrelated to each other (shuffled), while still possessing identical gene numbers, lengths, nucleotide compositions, and codon distributions compared to unshuffled melanin biosynthetic genes.

To detect homologous genes (paralogous or orthologous genes), we chose a conservative E-value of 10^−541^ as our significance threshold. From the Gaussian distribution of shuffled genes (Figure 8A), we determined that an E-value of 10^−5^ corresponds to a pairwise sequence alignment identity of 42%. In other words, any two genes possessing a pairwise sequence alignment identity ≥ 42% would be expected to occur in 1 out of 100,000 (1/10^−5^) randomly shuffled datasets by chance alone.

We detected 6 pairs of genes possessing pairwise sequence alignment identities ≥ 42% (E ≤ 10^−5^) (Figure 8A), corresponding to 11 individual genes. We constructed a maximum parsimony tree (Figure 8B). All genes-pairs possessing sequence alignment identities ≥ 42% (E ≤ 10^−5^) were found to belong to the same species (paralogs). No gene-pairs possessing sequence alignment identities ≥ 42% (E ≤ 10^−5^) were found to belong to different species (orthologs).

These results indicate that no orthologous genes exist for melanin biosynthesis across these four fungal strains. These results are expected as individual fungal strains are known to possess a large number of melanin-synthetic isozymes that convergently evolved to catalyze the same reactions^42^. It stands to reason that if colloidal melanin originated from a common ancestor, the descendant strains would retain the ancestral biochemical machinery for melanin synthesis, resulting in orthologs across the L-DOPA, DHN, and HGA pathways. The absence of detectable orthologs therefore indicates the absence of common ancestry, corroborating the alternate hypothesis of the convergent evolution of colloidal melanin established through melanin inhibitor assays (Figure 4).

## Discussion

In this study, we discovered and characterized a new class of water-soluble melanins (colloidal melanin) secreted by four strains of Ascomycetous fungi: *Gliomastix polychroma* (VIG 6138 / NFCCI 6138), *Stachybotrys chlorohalonatus* (VIG 2586). *Curvularia sp*. (VIG 2862), and *Pseudopithomyces sp*. (VIG 728). Our exploration was prompted by the recognition that soluble melanins represent rare exceptions within the predominantly insoluble melanin family. We have probed the polymer classes’ nature using spectroscopic, mass-spectrometric, electron microscopic, genomic, phylogenetic, and biochemical techniques. We present our experimental results as 9 inferences (Figure 9), explained below.

**Figure 9.**
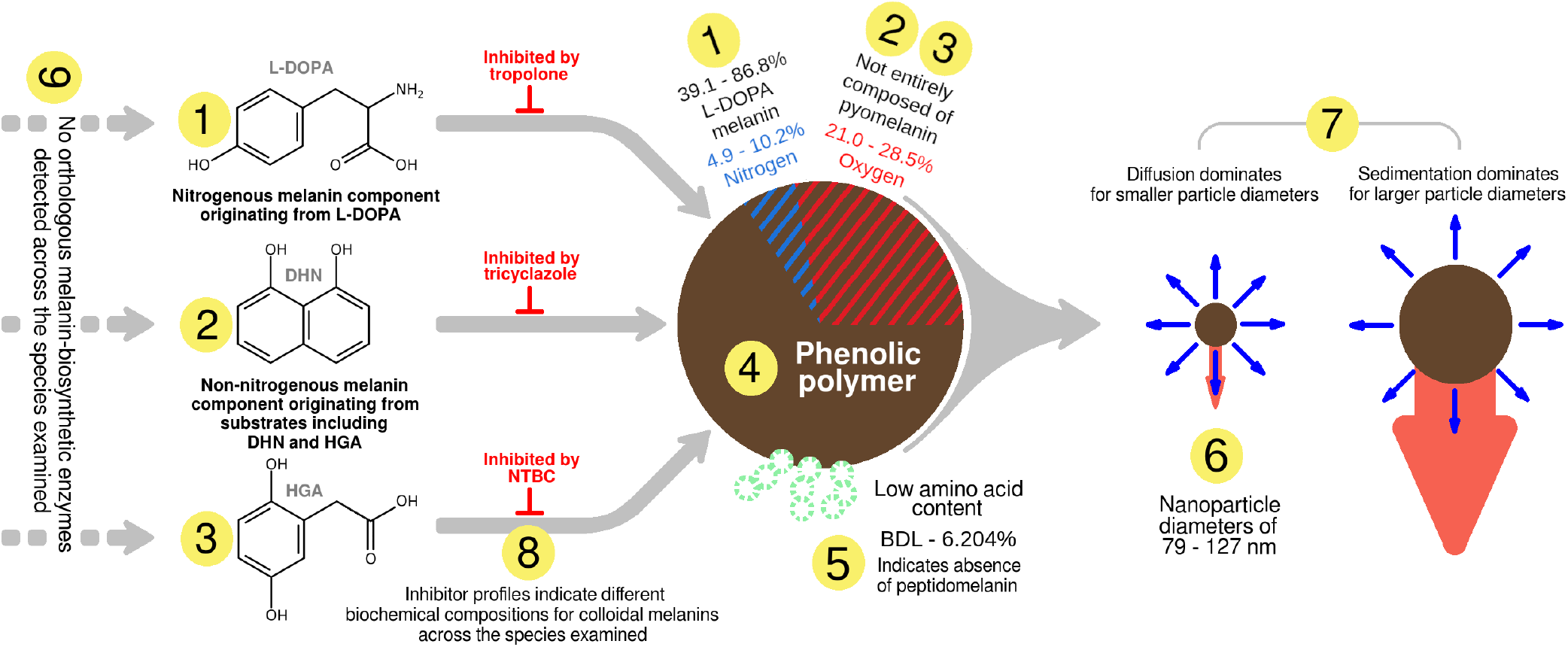
A summary of the inferences pertaining to the structure, biochemical composition, and biosynthesis of colloidal melanin. A detailed explanation of inferences 1-9 is provided in the text.

### Inference 1

Colloidal melanin possesses an L-DOPA component. This was inferred from the presence of nitrogen in the polymer, detected via SEM-EDS (Figure 5A,B, Table 3), the inhibition of soluble melanin synthesis using tropolone (Figure 4), and the identification of genes involved in the L-DOPA pathway from the organisms’ whole genome sequences (Table 5). SEM-EDS spectra (Table 3) revealed that colloidal melanin polymers possess a nitrogen content of 5.0 - 10.2% (wt%), corresponding to an L-DOPA content of 39.1 to 85.8% (wt%) (Figure 5B) after subtracting the nitrogen contribution of amino acids (Table 2). Additionally, the low nitrogen content indicates that colloidal melanin is not solubilized via nitrogenous functional groups.

### Inference 2,3

Colloidal melanin possesses a non-nitrogenous melanin component, with evidence for DHN and HGA precursors. This was inferred from a lower than expected nitrogen signal in the polymer (detected via SEM-EDS, Figure 5B), had the polymer been composed entirely of L-DOPA. The inhibition of colloidal melanin synthesis using tricyclazole and NTBC (Figure 4), along with the identification of genes responsible for DHN and HGA melanin synthesis (Table 5), further corroborated this inference. Colloidal melanin polymers possess an oxygen content of 21.0 to 28.5% (Figure 5C, Table 3), compared to the 44.8% oxygen content for synthetic pyomelanin. Although it was not possible to quantify the pyomelanin content due to the presence of multiple oxygen-containing monomers, these results nevertheless show that pyomelanin, although present, is not a major component of colloidal melanin. Additionally, the low oxygen content indicates that colloidal melanin is not solubilized via oxygenous functional groups.

### Inference 4

Colloidal melanin is phenolic in composition. This was expected as all natural melanins are derived from phenolic precursors, and was confirmed using FTIR (Figure 3C).

### Inference 5

Colloidal melanin is not composed of peptidomelanin. Colloidal melanin possesses an amino acid content of 0 to 6.204% (Table 2), significantly lower than the 22.98% amino acid content of peptidomelanin.

### Inference 6

Colloidal melanin is composed of nanoparticles with mean diameters ranging from 79 to 127 nm (Figure 7E, Table 4).

### Inference 7

The small diameters of colloidal melanin enable its apparent solubilization via colloidal dispersion. This was demonstrated using both theoretical (Table 4) and empirical (Figure 6) frameworks.

### Inference 8

Melanin inhibition experiments (Figure 4) indicate that colloidal melanin polymers secreted by different fungal strains are synthesized through distinct combinations of biosynthetic pathways, rather than through a single conserved pathway. This finding suggests that colloidal melanin evolved convergently.

### Inference 9

provides further support for inference 8. The absence of orthologous melanin biosynthetic enzymes across the four strains examined indicates that each lineage independently recruited its own biosynthetic machinery to produce colloidal melanin, rather than inheriting a shared colloidal melanin pathway from a common ancestor. We propose that convergence toward colloidal melanin is evolutionarily plausible because it is synthesized along the same biochemical pathways as known insoluble melanin polymers (L-DOPA, DHN, and HGA melanin). Therefore, the transition from an insoluble to a colloidal form appears to require only a biophysical change (reduction in particle diameter) rather than a biochemical one (creating a new biosynthetic pathway). We hypothesize that simple quantitative shifts in the *in vivo* reaction rates or reaction conditions could convert a parent insoluble melanin polymer into its colloidal counterpart.

In summary, we have addressed the question of what makes colloidal melanin soluble, characterizing its biophysical and biochemical properties. The question of why colloidal melanin is produced, however, remains unanswered. Future work will focus on understanding the ecological role of colloidal melanin and the evolutionary advantages it confers to these fungal strains.

## Methods

### Culture isolation and melanin purification

For melanin purification, all strains were cultured in Czapek Dox broth containing 0.5% tyrosine and incubated at 26 ± 2^°^C under light for 30 days. The culture filtrate was centrifuged (9500 rpm, 5^°^C) to remove mycelial biomass and the supernatant was treated with 100 mM KCl–HCl buffer (pH 2.0) to precipitate pigment. The precipitate was resolublized in 100 mM tris (at pH 8.0), purified by filtration using a 0.2 *µ*M polyvinylidene fluoride (PVDF) syringe filter, re-precipitated in 100 mM KCl–HCl buffer (pH 2.0), centrifuged (9500 rpm, 5^°^C), and the pellet was dried at 80 ^°^C in the dark for 24 hours, forming a solid powder. For experimental use, the pigment was dissolved in DMSO and dialyzed twice against 1L of deionized water over 48 hours.

### Chemical synthesis of melanins

#### For spectral characterization of L-DOPA melanin

5 mL of 2 mg/mL L-DOPA in 100 mM tris at pH 9.0 added in the presence of 1mM CuSO_4_(catalyst) was polymerised. We have previously synthesized L-DOPA melanin using a similar approach^23,25^.

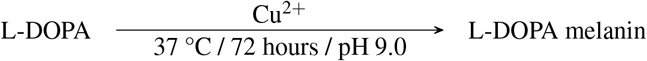

The reaction was incubated at 37 °C for 72 hours, then dialyzed through a 3 kDa dialysis membrane twice against 1 L of deionized water over 48 hours.

#### Synthesis of insoluble L-DOPA melanin nanoparticles

40 mg of L-DOPA in 10 mL of DMSO was incubated at 80°C for 7 days in presence of 1mM CuSO_4_ (catalyst).

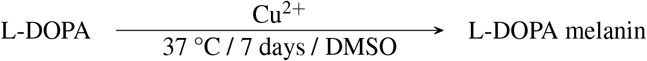

#### Synthesis of soluble L-DOPA melanin nanoparticles

100 *µ*L of 50 mg/mL KMnO_4_ solution was added to 10 mg L-DOPA in 1 mL of deionised water and then incubated for 10 min at 30°C. 10 mg of DTT was added to stop the reaction immediately after 10 minutes

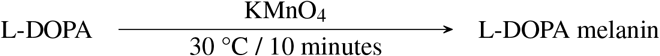

The reaction was centrifuged at 7000 rmp for 10 min to remove the byproducts (K_2_O and MnO_2_).

#### Synthesis of pyomelanin from homogentisic acid

100 mg of homogentisic acid was added to 5 mL of 100 mM tris at pH 10 with 50 *µ*L of 100 mM CuSO_4_.

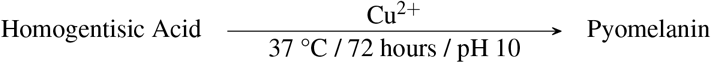

The reaction was incubated at 37 °C for 72 hours, then dialyzed through a 3 kDa dialysis membrane against 2 × 1 L of deionized water over 48 hours. Dialysed sample was then dried at 80 °C for 24 hours.

### Scanning electron microscopy - energy dispersive X-ray spectroscopy (SEM-EDS)

SEM/SEM-EDS was carried out using a FEI Quanta 200 instrument (Field Electron and Ion Company) at Icon Laboratories Pvt. Ltd., Mumbai. Samples were examined at an accelerating voltage of 20 kV under low-vacuum conditions with a chamber pressure of 65 Pa.

### UV – visible spectroscopy

UV–vis spectra were recorded using a Shimadzu UV 1900i spectrophotometer. Absorbance was measured from 200 to 1000 nm at a resolution of 0.5 nm, with five scans collected per sample and averaged. All spectra were obtained in deionized water.

### FTIR spectroscopy

FTIR spectra of all melanin samples were obtained using a Bruker Alpha II Compact FTIR spectrometer. Measurements were performed in ATR transmission mode across 3500–500 cm^−1^ with a resolution of 2 cm^−1^, averaging 16 scans per sample. Prior to analysis, all melanin samples were dialyzed against deionized water, dried at 80^°^C overnight to a powder form.

### Acid hydrolysis and amino acid analysis

The protocol provided below was used to separately quantify the amino acid content of each of the four colloidal melanin samples described in this study.

4 tubes containing 500 *µ*L (2 mL in total) of colloidal melanin (from a single fungal species) at a concentration of 1.2 mg/ml were acid-hydrolyzed by adding 500 *µ*L of 12 N HCl containing 1% phenol, for a total volume of 1 mL per tube (4 mL in total). The concentration of colloidal melanin per tube is therefore reduced to 0.6 mg/mL.

Samples underwent acid hydrolysis for 24 hours at 80^°^C under a nitrogen atmosphere. Following hydrolysis, the insoluble melanin fraction was pelleted from each tube by centrifugation (7,800 rpm / 10 mins), and the resulting supernatants were collected. Each supernatant was then dried at 80^°^C to remove residual HCl, leaving behind a residue containing amino acids. The residue in each tube was resuspended in 1 mL of deionized water (4 mL total). The resuspended contents of two tubes were pooled together to obtain 2 samples (replicates) of 2 mL volume each (4 mL in total).

All samples were sent to Om Diagnostics (Bangalore, India) to obtain the amino acid composition using the amino acid profile

1. test. Raw data from this test for all colloidal melanin samples is provided in **Dataset_S3:LC-MS_data**. Note that as this is a clinical test, a clinical interpretation is included with the raw data, which can be disregarded. Calculations used to convert the raw data into the values presented in Table 2 are also provided for all replicates in the supplementary information (**Dataset_S3:LC-MS_data**).

### Melanin synthesis inhibition assays

Czapek Dox medium (10 mL) was supplemented with either the DHN melanin synthesis inhibitor tricyclazole (Dow Agro Sciences; 100 *µ*g/mL), the L-DOPA melanin synthesis inhibitor tropolone (Sigma Aldrich; 2 mM), or the pyomelanin (HGA melanin) synthesis inhibitor NTBC (2-(2-nitro-4-trifluoromethylbenzoyl)1,3-cyclohexanedione) (Merck; 1 mM). Each medium was inoculated with a 5 mm diameter mycelial plug excised from a colony grown on Czapek Dox agar and incubated for 10 days at 26 *±* 2 ^°^C. Medium without inhibitor served as the control.

### Transmission electron microscopy (TEM)

Colloidal melanin samples were diluted with 4.0 mL of deionized water, and 0.2 *µ*L of the suspension was placed onto a carbon coated copper grid (200 mesh). The grids were air dried at room temperature, negatively stained with 2% phosphotungstic acid (PTA) for 20 seconds, and excess stain was removed with filter paper. Observations were carried out using a TECNAI G2 Spirit BioTWIN transmission electron microscope (FEI, Netherlands) fitted with a tungsten (W) filament and operated at 100–120 kV. Images were acquired with an Olympus Soft Imaging Solutions VELETA CCD camera and analyzed using Tecnai Imaging and Analysis software. The instrument allowed magnifications up to 350,000*×*.

### Whole genome sequencing

Whole genome sequencing was performed by Nucleome Informatics Pvt. Ltd. (Hyderabad, India). Genomic DNA was extracted from mature fungal biomass using the DNeasy PowerSoil Pro Kit (Qiagen) and was assessed for quality using spectrophotometric and fluorometric methods. Sequencing libraries were prepared using the KAPA HyperPlus Kit (Roche) and were sequenced on an Illumina NovaSeq 6000 platform using an S4 flow cell.

Raw sequencing reads were quality filtered and used for *de novo* genome assembly using SPAdes(v3.15.5)^43^ assembler. Assembly quality and completeness were evaluated using standard assembly metrics and Benchmarking Universal Single-Copy Orthologs (BUSCO)^44^ with a fungal lineage dataset. Gene prediction was performed using GeneMark-ES, and functional annotation was conducted using EggNOG-mapper^45^and KEGG pathway analysis.

Annotated genes were screened to identify enzymes involved in fungal melanin biosynthesis, including the L-DOPA melanin, DHN melanin, and HGA melanin pathways.

### Phylogenetics

Phylogenetic analysis for melanin biosynthetic genes extracted from four fungal strains was performed using MEGA6^46^ (Figure 8B). All gene sequences are provided in the supplementary information (**Dataset_S5:annotated_gene_sequences**). Gene sequences were initially aligned using MUSCLE^47^. Alignments were generated with a gap open penalty of -400.00 and a gap extension penalty of 0.00. The maximum memory allocation was set to 2048 MB and the maximum number of iterations to 16. UPGMA was used as the clustering method for both the first and subsequent iterations, and the minimum diagonal length (lambda) was set to 24. All other parameters were left at their default values.

Phylogenetic analysis was conducted in MEGA6 using the Maximum Parsimony (MP) method. The MP tree was obtained using the Subtree-Pruning-Regrafting (SPR) search method with 10 initial trees obtained by random addition (search level 1), retaining a maximum of 100 most parsimonious trees. Nucleotide sequences were analyzed with all sites included (gaps and missing data were not excluded). Branch support was assessed using the bootstrap method with 100 replications. Analyses were run using 3 threads.

### Statistical Analysis

All statistical analyses were performed using R version 4.5.1 (2025-06-13) - “Great Square Root”.

#### Use of a regression model to classify melanin nanoparticle solubility (Figure 6E)

Particle diameters of soluble and insoluble L-DOPA melanin nanoparticles were compared using binary logistic regression using the glm() function in R, with family = “binomial”. Solubility (soluble = 1, insoluble = 0) was modeled as a function of particle diameter using a generalized linear model with a binomial error distribution and logit link function. The model was fitted using individual nanoparticle diameter measurements extracted from TEM data (**Dataset_S2:TEM_data**). Predicted probabilities of solubility across the observed diameter range were generated along with 95% confidence intervals (fitted value *±* 1.96 SE on the link scale, back-transformed via the inverse logit function).

#### One way ANOVAs and Post-hoc pairwise comparisons (Figures 5A,C and 7E,F)

Differences among cohorts were assessed using a one-way analysis of variance (ANOVA; aov). Post-hoc pairwise comparisons were performed using one-tailed Welch’s t-tests (t.test, unequal variances, not pooled). The resulting p-values were corrected for multiple comparisons using the Holm method (p.adjust, method = “holm”).

For Figure 7E,F: pairwise comparisons against insoluble L-DOPA melanin were performed using alternative “greater” (red p-values and arrows), in order to answer the specific question “are soluble melanin nanoparticles significantly *smaller* than insoluble L-DOPA melanin nanoparticles?” Likewise, pairwise comparisons against soluble L-DOPA melanin were performed using the alternative “less” (blue p-values and arroes), in order to answer the specific question “are fungal colloidal melanin nanoparticles *larger* than soluble L-DOPA melanin nanoparticles?”

#### Pairwise comparisons for amino acid content (Table 2)

Statistical significance testing was performed using an independent two-sample Student’s *t*-test, assuming equal variances between groups. For each comparison, the mean 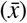, standard deviation (*SD*), and sample size (*n*) of each group were used to calculate a pooled variance 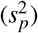 as:

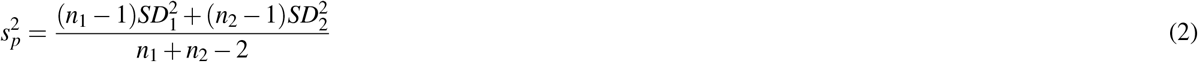

The pooled standard deviation (*s*_*p*_) was then used to compute the *t*-statistic as:

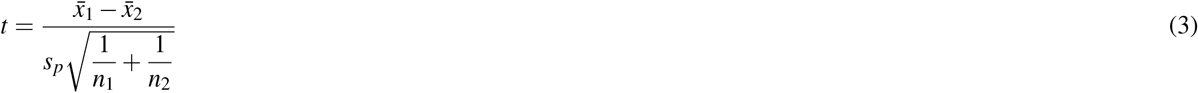

with degrees of freedom equal to *n*_1_ + *n*_2_ − 2. Two-tailed *p*-values were derived from the *t*-distribution using the calculated *t*-statistic and degrees of freedom. Comparisons were computed using a custom function in R (**Dataset_S3:LC-MS_data**). Amino acids reported as below the limit of detection (BDL) were excluded from statistical comparison, as no variance estimate was available for these groups.

#### E value estimation (Figure 8A)

To assess the significance of pairwise sequence identity among melanin biosynthetic genes, a null distribution was generated by shuffling each sequence and recalculating pairwise identity matrices (ClustalW pairwise alignment) for both the unshuffled and shuffled sequence sets. A Gaussian distribution was fitted to the sequence identity values of the shuffled (null) dataset using maximum likelihood estimation (fitdist, package fitdistrplus), yielding a mean and standard deviation describing the expected distribution of sequence identity arising by chance.

An E-value was calculated for a given sequence identity value as the probability density of that value under the fitted null distribution (dnorm), multiplied by the number of pairwise comparisons in the shuffled dataset (analogous to the expectation value used in BLAST searches, representing the expected number of chance matches at or above a given identity under the null model). The sequence identity threshold corresponding to E = 10^−5^ was determined by numerically scanning identity values between the null distribution mean and 50% in increments of 0.001% and identifying the value whose computed E-value was closest to the target.

## Acknowledgments

Authors Govinda Rajulu MB, Thirunavukkarasu N, Ravishankar JP, and Suryanarayanan TS thank Swami Dhyanagamyananda, Secretary, Ramakrishna Mission Vidyapith, Chennai, for providing facilities. The authors thank Mr. Kiran Rambhau Bhotkar and Mrs. Sunita Samgir (Icon Labs Pvt. Ltd., Mumbai) for their expert assistance with scanning electron microscopy. The authors also acknowledge Mr. Venkatesh T. and Ms. Bhagyashree Thorat (Icon Labs Pvt. Ltd., Mumbai) for their technical support with transmission electron microscopy. The authors thank Connor Grigg for the useful discussions.

## Author contribution statement

Authors Senjuti Sarkar, Ananya Krishnaswamy, Govinda Rajulu MB, Pema Lhamu, Bhavana Chamraj, Deepika Rai, and Rakshita S. Kolipakala performed all experiments. Authors Huma Sulthana and Judy Jays collected FTIR spectra. Authors Ananya Krishnaswamy. and Deepesh Nagarajan performed data analysis and statistical analyses. Authors Shiwali Rana and Sanjay K Singh performed Sanger Sequencing and phylogenetic analysis for strain identification. Authors Ananya Krishnaswamy. and Pushya Pradeep analyzed and annotated whole genome sequences. Authors Thirunavukkarasu N, Ravishankar JP, Suryanarayanan TS, and Deepesh Nagarajan conceived the study and designed all experiments. All authors participated in writing and reviewing the manuscript before submission.

## Supplementary information

All supplementary information has been made publicly available on Figshare (DOI: 10.6084/m9.figshare.33088712): https://doi.org/10.6084/m9.figshare.33088712.

**Dataset S1:SEM_data**: Supplementary data for Figure 2. This dataset contains all SEM micrographs collected for colonies of *Gliomastix polychroma* (VIG 6138 / NFCCI 6138), *Stachybotrys chlorohalonatus* (VIG 2586), *Curvularia sp*. (VIG 2862), and *Pseudopithomyces sp*. (VIG 728). All cultures were grown on Sabouraud dextrose agar for 30 days. Cultures were refrigerated prior to environmental SEM visualization.

**Dataset S2:TEM_data**: Supplementary data for Figure 6 and Figure 7. This dataset contains TEM micrographs depicting individual nanoparticles for the following melanin classes: L-DOPA melanin (soluble), L-DOPA melanin (insoluble), colloidal melanin from *Gliomastix polychroma* (VIG 6138 / NFCCI 6138), colloidal melanin from *Stachybotrys chlorohalonatus* (VIG 2586), colloidal melanin from *Curvularia sp*. (VIG 2862), and colloidal melanin from textitPseudopithomyces sp. (VIG 728). R and bash scripts for statistical analysis and data visualization are also provided.

**Dataset S3:LC-MS_data** Supplementary data for Table 2. This dataset contains LC-MS reports for acid-hydrolyzed melanin obtained from *Gliomastix polychroma* (VIG 6138 / NFCCI 6138), *Stachybotrys chlorohalonatus* (VIG 2586), *Curvularia sp*. (VIG 2862), and *Pseudopithomyces sp*. (VIG 728) are provided. Spreadsheets and R scripts for data analysis and statistical analysis are also provided.

**Dataset S4:SEM-EDS_data** Supplementary data for Figure 5 and Table 3 are provided. This dataset contains SEM-EDS reports containing the elemental compositions of colloidal melanin obtained from *Gliomastix polychroma* (VIG 6138 / NFCCI 6138), *Stachybotrys chlorohalonatus* (VIG 2586), *Curvularia sp*. (VIG 2862), and *Pseudopithomyces sp*. (VIG 728). Spreadsheets, R and bash scripts for statistical analysis and data visualization are also provided.

**Dataset S5:annotated_gene_sequences**: Supplementary data for Table 5, Figure 8, and Supplementary Tables S1-S4. This dataset contains genes identified from whole-genome sequencing (WGS) and annotated as putatively involved in melanin biosynthetic pathways in *Gliomastix polychroma* (VIG 6138 / NFCCI 6138), *Stachybotrys chlorohalonatus* (VIG 2586), *Curvularia sp*. (VIG 2862), and *Pseudopithomyces sp*. (VIG 728). Functional annotations were assigned based on EggNOG-marker and KEGG pathway analysis predictions and homology to known L-DOPA, DHN, and HGA melanin biosynthetic enzymes. Gene data is provided in .fasta format. Scripts in Python, R, and bash are provided for data analysis, statistical analysis, and data visualization. Raw data used to construct the phylogenetic tree in Figure 8B is provided as a .meg (MEGA6) file.

**Text S1:** Multilocus molecular identification and phylogenetic analysis for *Gliomastix polychroma* (VIG 6138 / NFCCI 6138).

**Text S2:** Species identification of *Stachybotrys chlorohalonatus* (VIG 2586).

**Text S3:** ITS sequences for all strains examined in this study: *Gliomastix polychroma* (VIG 6138 / NFCCI 6138), *Stachybotrys chlorohalonatus* (VIG 2586), *Curvularia sp*. (VIG 2862), and *Pseudopithomyces sp*. (VIG 728).

**Table S1:** Supplementary data for Table 5. Genes from *Gliomastix polychroma* (VIG 6138 / NFCCI 6138) identified from whole-genome sequencing (WGS) and annotated as putatively involved in melanin biosynthetic pathways. Functional annotations were assigned based on EggNOG-mapper and KEGG pathway analysis predictions and homology to known L-DOPA, DHN, and HGA melanin biosynthetic enzymes.

**Table S2:** Supplementary data for Table 5. Genes from *Stachybotrys chlorohalonatus* (VIG 2586) identified from whole-genome sequencing (WGS) and annotated as putatively involved in melanin biosynthetic pathways. Functional annotations were assigned based on EggNOG-mapper and KEGG pathway analysis predictions and homology to known L-DOPA, DHN, and HGA melanin biosynthetic enzymes.

**Table S3:** Supplementary data for Table 5. Genes from *Curvularia sp*. (VIG 2862) identified from whole-genome sequencing (WGS) and annotated as putatively involved in melanin biosynthetic pathways. Functional annotations were assigned based on EggNOG-mapper and KEGG pathway analysis predictions and homology to known L-DOPA, DHN, and HGA melanin biosynthetic enzymes.

**Table S4:** Supplementary data for Table 5. Genes from *Pseudopithomyces sp*. (VIG 728) identified from whole-genome sequencing (WGS) and annotated as putatively involved in melanin biosynthetic pathways. Functional annotations were assigned based on EggNOG-mapper and KEGG pathway analysis predictions and homology to known L-DOPA, DHN, and HGA melanin biosynthetic enzymes.

## Data availability

The following depositions have been made for genetic and genomic data from *Gliomastix polychroma* (VIG 6138 / NFCCI 6138). Whole-genome sequence: GenBank accession ID: JBSVRS000000000.1. Internal transcribed spacer (ITS): GenBank accession ID: PX114506. Large subunit ribosomal RNA gene (LSU): GenBank accession ID: PX114507. Translation elongation factor 1-alpha (*EF-1α*): GenBank accession ID: PX124606. It should be noted that our strain of *Gliomastix polychroma* was previously designated *Acremonium polychromum*. The name was corrected based on a recent taxonomic revision^48^. Some sequence data may contain the previous nomenclature.

The following depositions have been made for genetic and genomic data from *Stachybotrys chlorohalonatus* (VIG 2586). Whole-genome sequence: GenBank accession ID: JBYLTQ000000000. Internal transcribed spacer (ITS): GenBank accession ID: PZ476833.

The whole-genome sequence for *Curvularia sp*. (VIG 2862) has been deposited under GenBank accession ID: JBZEGM000000000.

The whole-genome sequence for *Pseudopithomyces sp*. (VIG 728) has been deposited under GenBank accession ID: JBZEGL000000000.

